# Molecular dynamics investigation of the influenza hemagglutinin conformational changes in acidic pH

**DOI:** 10.1101/2024.07.07.602399

**Authors:** Shadi A Badiee, Vivek Govind Kumar, Mahmoud Moradi

## Abstract

The surface protein hemagglutinin (HA) of the influenza virus plays a pivotal role in facilitating viral infection by binding to sialic acid receptors on host cells. Its conformational state is pH-sensitive, impacting its receptor-binding ability and evasion of the host immune response. In this study, we conducted extensive equilibrium microsecond-level all-atom molecular dynamics (MD) simulations of the HA protein to explore the influence of low pH on its conformational dynamics. Specifically, we investigated the impact of protonation on conserved histidine residues (His106_2_) located in the hinge region of HA2. Our analysis encompassed comparisons between non-protonated (NP), partially protonated (1P, 2P), and fully-protonated (3P) conditions. Our findings reveal substantial pH-dependent conformational alterations in the HA protein, affecting its receptor-binding capability and immune evasion potential. Notably, the non-protonated form exhibits greater stability compared to protonated states. Conformational shifts in the central helices of HA2 involve outward movement, counterclockwise rotation of protonated helices, and fusion peptide release in protonated systems. Disruption of hydrogen bonds between the fusion peptide and central helices of HA2 drives this release. Moreover, HA1 separation is more likely in the fully-protonated system (3P) compared to non-protonated systems (NP), underscoring the influence of protonation. These insights shed light on influenza virus infection mechanisms and may inform the development of novel antiviral drugs targeting HA protein and pH-responsive drug delivery systems for influenza.

## Introduction

Thousands of people suffer from influenza infections every year, but there is not any fundamental cure for them. Therefore, it is necessary to thoroughly understand the infection process and the critical keys to preventing infections. An influenza virus can be classified into four sub-types: A, B, C, and D. Those belonging to groups A and B contain two membrane-embedded glycoproteins named hemagglutinin (HA) and neuraminidase (NA), while those belonging to groups C and D contain only one surface glycoprotein, known as hemagglutinin-esterase fusion (HEF).^1–4^ Influenza A and B viruses are responsible for most human infections and are responsible for seasonal epidemics and occasional pandemics. The influenza virus uses glycoproteins on its surface to enter host cells,^3^ and understanding the structural features of these proteins that make them vulnerable to inhibitors is important for the development of anti-viral drugs. As part of the fusion process, viral hemagglutinin (HA) plays an important role.^5^ Influenza viruses have been very widely studied and are well characterized as a global health concern. However, the atomistic details of influenza HA-mediated membrane fusion are still poorly understood.^6,7^

Hemagglutinin (HA) plays a key role in the virus’s ability to infect host cells.^8–11^ Hemagglutinin binds to sialic acid receptors on the surface of viral membrane, which allows the virus to enter and replicate within host cells.^12^ Hemagglutinin (HA) protein is synthesized as a precursor protein, known as HA0, which is then cleaved by a cellular protease to generate the mature HA protein.^10,13,14^ The Hemagglutinin (HA) has a trimeric structure composed of three identical monomers, each monomer is composed of two main subunits, HA1 and HA2 (Fig. 1). These two subunits are covalently held together by disulfide bond ^15^ and form the functional unit of the protein. HA1 forms the globular head of the protein which is responsible for binding to host cell receptors and initiating the viral entry into the host cell.^10^ In comparison to HA1, HA2 is smaller. It contains the fusion machinery and is responsible for anchoring the HA protein to the viral envelope. It is composed of the stem domain which is important for the stability of the protein and the interactions between the HA and the other viral proteins .^10,16,17^ The HA2 subunit comprises the fusion peptide (FP, residues 1–20), two *β*-strands (TBS, residues 21–37), helix A (residues 38–54), B loop (residues 55–75), coiled coil (residues 76–104), hinge region (residues 105–129), and ectodomain and transmembrane domain (TMD, residues 130–175).^18^ Additionally, domains in HA1 include the fusion (F*^′^*), vestigial esterase (VE), and receptor-binding domain (RBD).^19,20^ The RBD contains the receptor-binding site (RBS), which is made up of four secondary structural elements: the 130-loop, the 150-loop, the 190-helix, and the 220-loop.^21^ The VE domain encompasses the 30-loop, which consists of HA1 residues 22–37.^22^ Throughout this paper, subscript 1 will be used for residues belonging to HA1, while subscript 2 will denote residues of HA2.

**Fig. 1.**
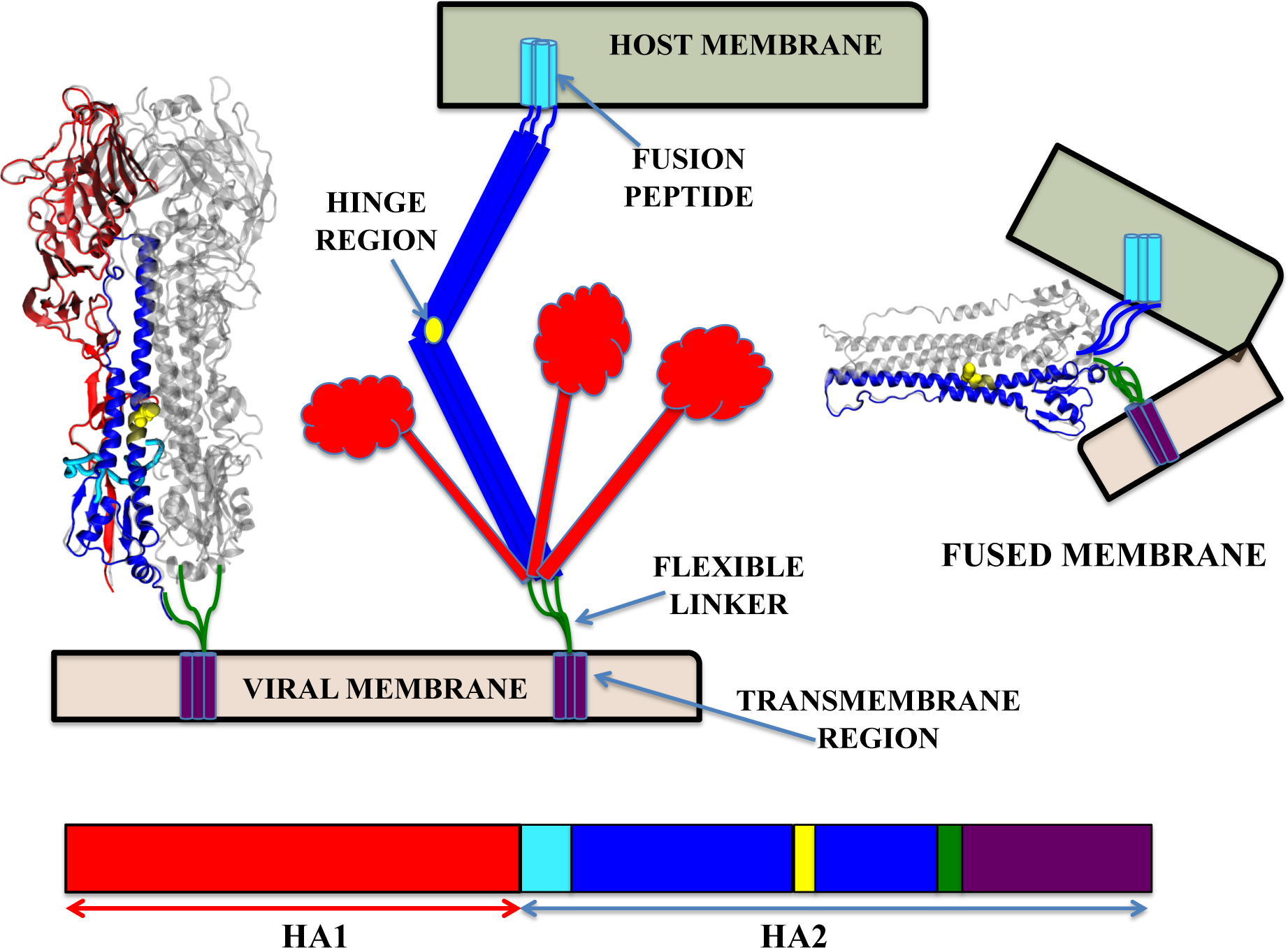
Schematic representation of Influenza HA protein-mediated membrane fusion: At low pH, the HA1 domains dissociate from the HA2 domains, exposing the fusion peptide. Subsequent conformational changes in HA2 facilitate fusion between the host membrane and viral envelope.

In response to acidic conditions or low pH environment, hemagglutinin (HA), undergoes extensive conformational changes that allow the N-terminal portion of the HA2 subunit, called FP, to be inserted into the membrane of the host endosome.^17,23,24^ HA1 is thought to move away from HA2, causing the exposure of a FP at the HA2 N-terminus .^13,14,22^ HA2 is then thought to bend at a hinge region, which eventually results in the lipid bilayers being pulled towards each other.^13,14^ The HA2 C-terminus consists of a transmembrane domain needs to span the bilayer in order to promote fusion.^25^ Alternatively, upon acidification, the major conformational change happen in HA2. The B-loop located between two helices in HA2 has a loop-to-helix transformation and forms an *α*-helix elongating the fusion peptide to the endosomal membrane^26–28^ (Fig. 1). Recent experimental evidence has shown that multiple HA trimers work cooperatively to cause membrane fusion.^29^ Studies have shown that the ideal pH for HA conformational changes is between 4.8 and 6.0.^15,30^ Histidine depends on degree of conservation and its position on HA protein was identified as a potential pH-sensing residue.^31^ Histidine is the only amino acid with a pKa value similar to the acidic pH required for HA-mediated membrane fusion. Therefore, conformational changes of proteins that require acidic environments can be triggered by protonating one or more histidines in their structures.^31–33^ Therefore, His184_1_ on HA1 and His106_2_, His111_2_, His142_2_, and His159_2_ on HA2 were suggested as conserved histidines that can potentially trigger the conformational change mediating membrane fusion in HA protein.^31^ HA’s fusion domain consists of all the proposed histidines except for His184_1_.^34^ Among these histidine residues, a detailed analysis has been conducted for His111_2_ and His106_2_ in subtype H3 and H5, respectively.^34–37^ Studies indicated that conformational changes on HA is increased through protonation of His106_2_/His111_2_, however, almost no effect was seen from the mutation on these residues.^34,36,37^ A number of virus groups with pH-dependent infectivity have provided evidence to support the histidine-switch hypothesis, which states that changes in pH can alter the conformational state of the virus proteins, leading to altered infectivity.^31,33,38–47^

In this study, we investigated the conformational rearrangements of influenza HA when it is exposed to an acidic environment. All-atom equilibrium MD simulations at the microsecond level have been used to study the effects of protonating one, two or three conserved HA2 hinge histidine residues on the conformational dynamics of HA. The data from our extensive equilibrium MD simulations reveal that the HA protein undergoes conformational changes even when a single histidine residue is protonated.

## Methods

Our simulations were conducted based on the crystal structure of the Group-2 HA (PDB entry: 5KUY)^48^ . In our simulations, we focused on the conserved histidine residue (His106_2_ which is His435 in PDB file), located in the hinge region, which was protonated. In the Results section, HA2 residues were renumbered for the sake of convenience and consistency with previous studies, despite having different numbers in the PDB file. For instance, residue His435 in the PDB file was renumbered as residue His106 in this study to align with established literature conventions. Throughout this paper, subscript 1 will be used for residues belonging to HA1, while subscript 2 will denote residues of HA2. To generate simulation inputs, we utilized the CHARMM GUI simulation input generator.^49,50^ Four different systems were constructed: a system with no protonation (NP), one with a single protonation (1P), another with two protonations (2P), and finally, a system with three protonations (3P). We conducted three replicates for each system. Each system comprised one protein, 0.15 M NaCl, and TIP3P water molecules.^51^ For systems with no protonation (NP) and one protonation (1P), each contained 138281 water molecules and had dimensions of 163 × 163 × 163 Å ^3^. Systems with two protonations (2P) and three protonations (3P) included 130646 water molecules and had dimensions of 160 × 160 × 160 Å ^3^. To describe all molecules accurately, we employed the CHARMM36 all-atom additive force field parameters.^52,53^ Preliminary MD simulations were conducted using NAMD 2.13^54^ before the production runs on Anton 2. During simulations, we employed a Langevin integrator with periodic boundary conditions at a temperature of 310 K, a collision frequency of 0.5/ps, and a time step of 2 fs. The pressure was maintained at 1 atm using the Nosé-Hoover Langevin piston method.^55,56^ For non-bonded interactions, a smoothed cutoff distance of 10*−*12 Å was applied, and long-range electrostatic interactions were calculated using the particle mesh Ewald (PME) method.^57^ Prior to equilibration, each system underwent energy minimization for 10,000 steps using the conjugate gradient algorithm.^58^ Equilibration consisted of an initial relaxation in the NVT ensemble followed by a 15 ns equilibrium simulation in the NPT ensemble. After equilibration, production runs were performed on Anton 2 for 2.4 *µs* with a time step of 2.4 fs. Two additional repetitions were carried out for all systems (NP, 1P, 2P, and 3P) on Anton2, referred to as repeat 2 and repeat 3 in the manuscript. To generate initial conformations for repeat 2, the original production run for each model was extended by 0.5 ns on TACC Stampede. Subsequently, the production runs were extended by an additional 0.5 ns to generate the initial conformations for repeat 3. In total, 28.8 *µs* of simulation data was generated across 12 systems, with 2.4 *µs* allocated to each of the systems. For the simulations on Anton2, pressure was maintained isotropically at 1 atm using the MTK barostat, and temperature was controlled at 310 K using the Nosé-Hoover thermostat.^55,56^ Long-range electrostatic interactions were computed using the FFT (fast Fourier transform) method implemented on Anton 2.^59^

For analysis, molecular snapshots, and visualization, we employed visual molecular dynamics (VMD).^60^ Hydrogen bonds and salt bridges were calculated using VMD plugins with cutoff distances of 4.0 Å and angles of 30*^◦^* for hydrogen bonds. Salt bridge distances were computed as the smallest distance between donor and acceptor atoms. Principal component analysis (PCA) was performed using PRODY software,^61^ considering only C*α*.

## Results and Discussion

The RMSD plots (Fig. 2) present the RMSD values for HA0 and its individual domains, HA1 and HA2, over the course of the simulations. The complexity and size of the HA0 protein pose a significant obstacle in obtaining convergence in our simulations, which last for a few microseconds. This indicates that additional simulation time is necessary to accurately capture the protein’s complete conformational changes and ensure reproducibility of the results. Despite the limited simulation time, we can still observe significant conformational changes and flexibility in both the NP and 3P systems. Notably, the 3P systems exhibit greater flexibility compared to the NP systems. This increased flexibility in the 3P systems suggests that protonation state significantly influences the dynamic behavior of the protein. To achieve true convergence and reproducibility, longer simulation times are required. However, within the constraints of the current simulation time, we have focused our analysis on specific regions of the protein. By doing so, we can provide a detailed examination of the conformational changes occurring in these regions. This targeted approach allows us to draw meaningful conclusions about the behavior of these key regions, even if a comprehensive understanding of the entire protein’s dynamics is not yet attainable.

**Fig. 2.**
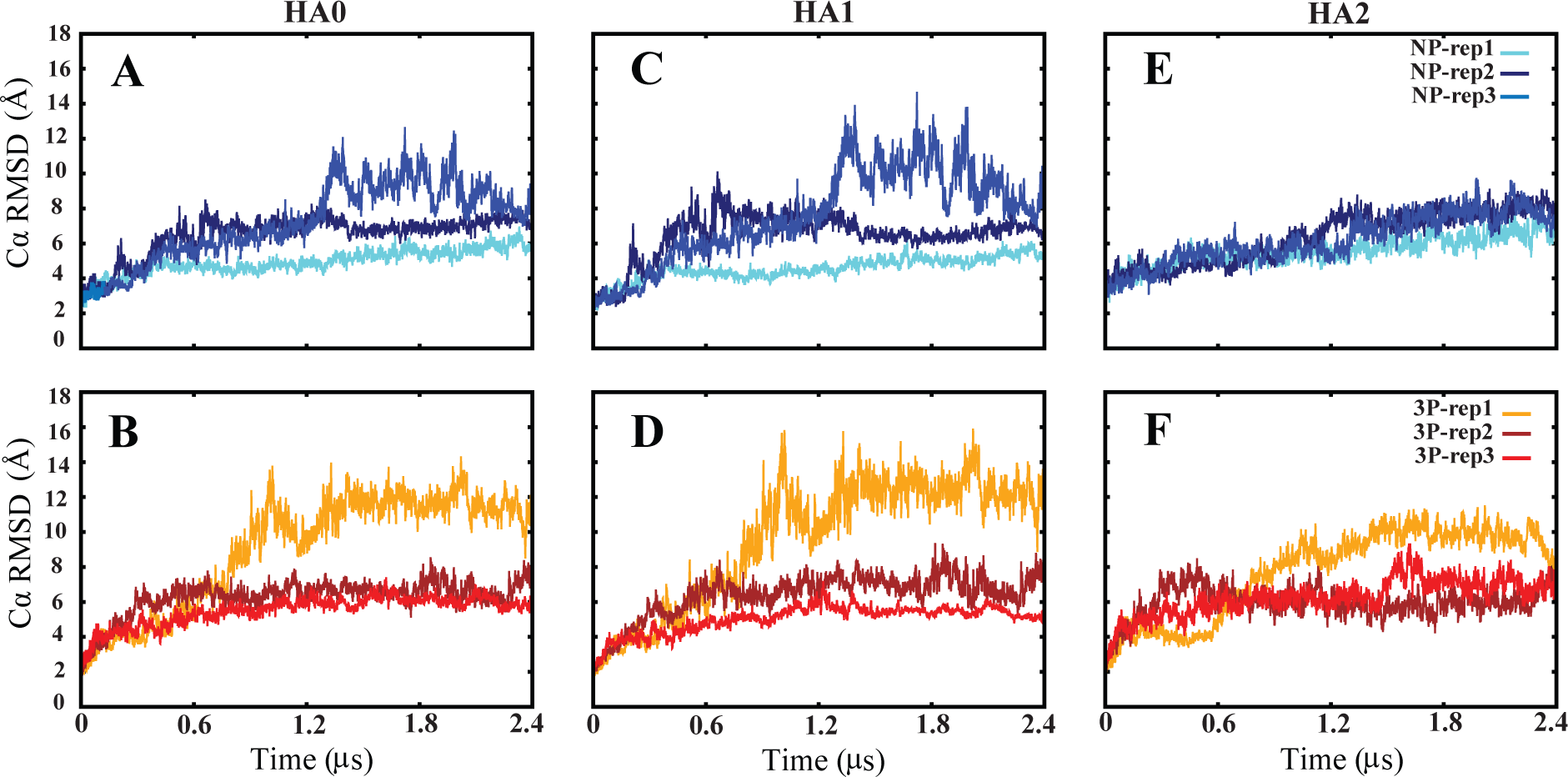
Time series of C*α* RMSD values for non-protonated (NP) and fully-protonated (3P) systems of HA0 (A & B), HA1 (C & D), and HA2 (E & F). Each NP and 3P system simulation was run three times for 2.4 *µs* each.

### Conformational dynamics of the S4 helix in fully-protonated system (3P) compared to non-protonated system (NP)

To determine the effect of protonated histidine and the stability of the S4 helix, which includes a conserved histidine, during our simulations, we measured its C*α* RMSD values (Fig. S1). In the NP systems, the RMSD stayed below 6 Å in all three sets (Fig. S1A). However, in the 3P systems, it went above 6 Å in two out of three repeats (Fig. S1B). We compared the last 1.2 *µs* of all repeats using RMSD distribution plots. The results indicate that the 3P systems exhibited higher flexibility, with RMSD values climbing up to 10 Å compared to 6 Å in the NP systems. This suggests that the protonation of histidine significantly impacts the stability of the S4 helix (Fig. 3).

**Fig. 3.**
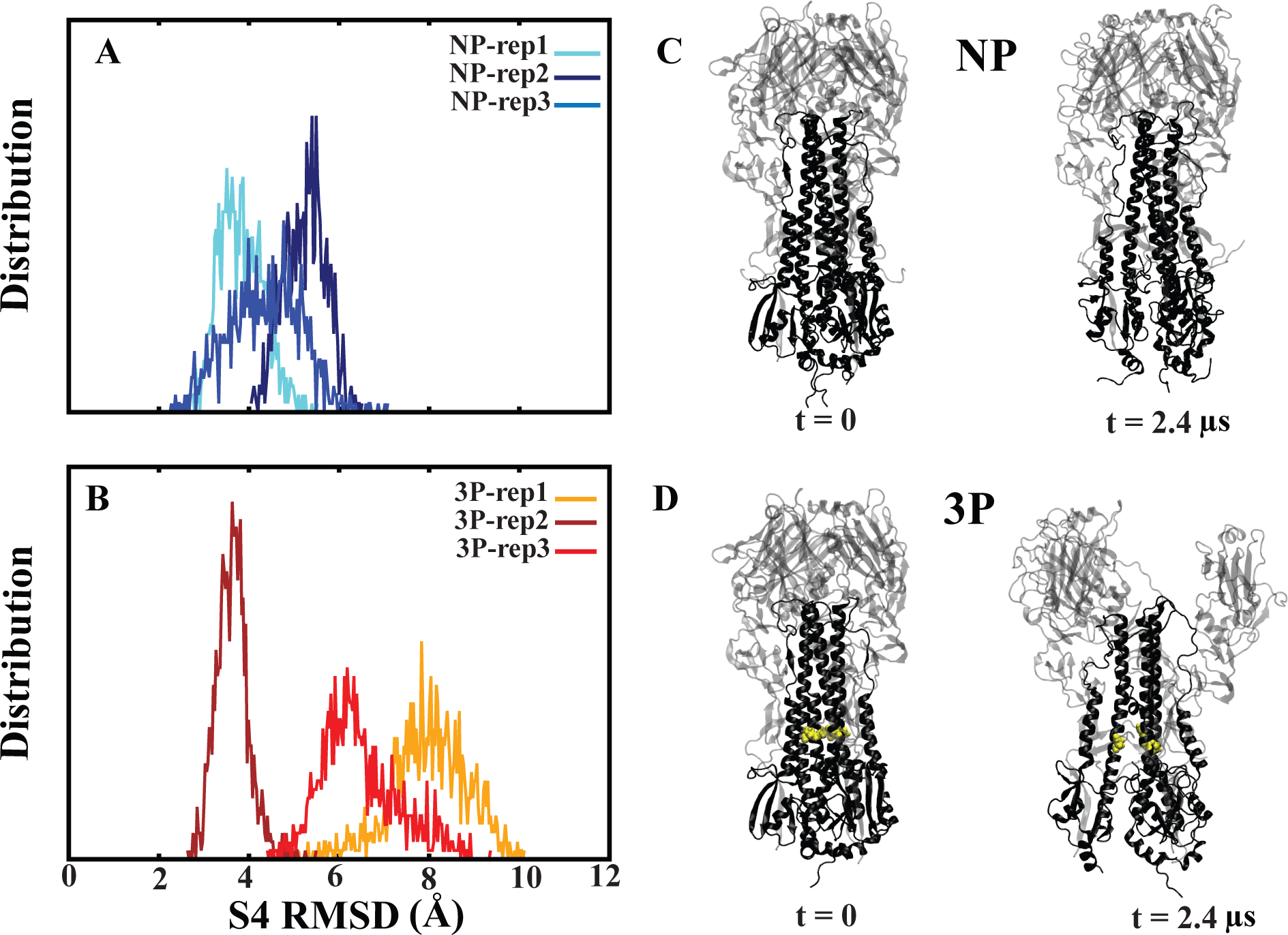
S4 RMSD distributions. Probability distribution of S4 helices RMSD values in non-protonated (NP) (A) and fully-protonated (3P) (B) systems during the final half of simulations (from 1.2 to 2.4 *µs*). (C & D) Cartoon representations of the non-protonated (NP) and fully-protonated (3P) system from the first and last frames of the simulation. The protonated histidine residues are shown with the yellow color.

We additionally quantified the quantity of water molecules within the lower region of the protein encompassing the three S4 domains (Fig. S1C & D). The findings reveal a greater abundance of water molecules in the system with three protonations (3P), particularly after 1.2 *µs* of simulation. Plotting the water molecule counts over the last 1.2 *µs*, distribution plots show that the 3P system’s peak is about 850 molecules, while the NP system’s peak is about 700 molecules (Fig. 4). The tilt angle analysis, focusing on S4’s tilt angle relative to the initial frame of HA2, indicates a gradual increase over the simulation duration in all repeats of 3P systems compared to NP systems. This reveals a contrasting trend, with the angle decreasing in the NP systems while consistently increasing in the 3P systems (Fig. S2). This suggests a dynamic shift where S4 approaches each other in the NP systems and moves away from HA2 in the 3P systems. Additionally, the average tilt angle analysis demonstrated that the tilt angle of all three S4 domains across the three repeats was higher in 3P compared to the overall average tilt angle in the NP system. Moreover, a more dispersed data distribution was observed in the 3P system, indicating a greater degree of opening in the fully-protonated environment (Fig. 5A).

**Fig. 4.**
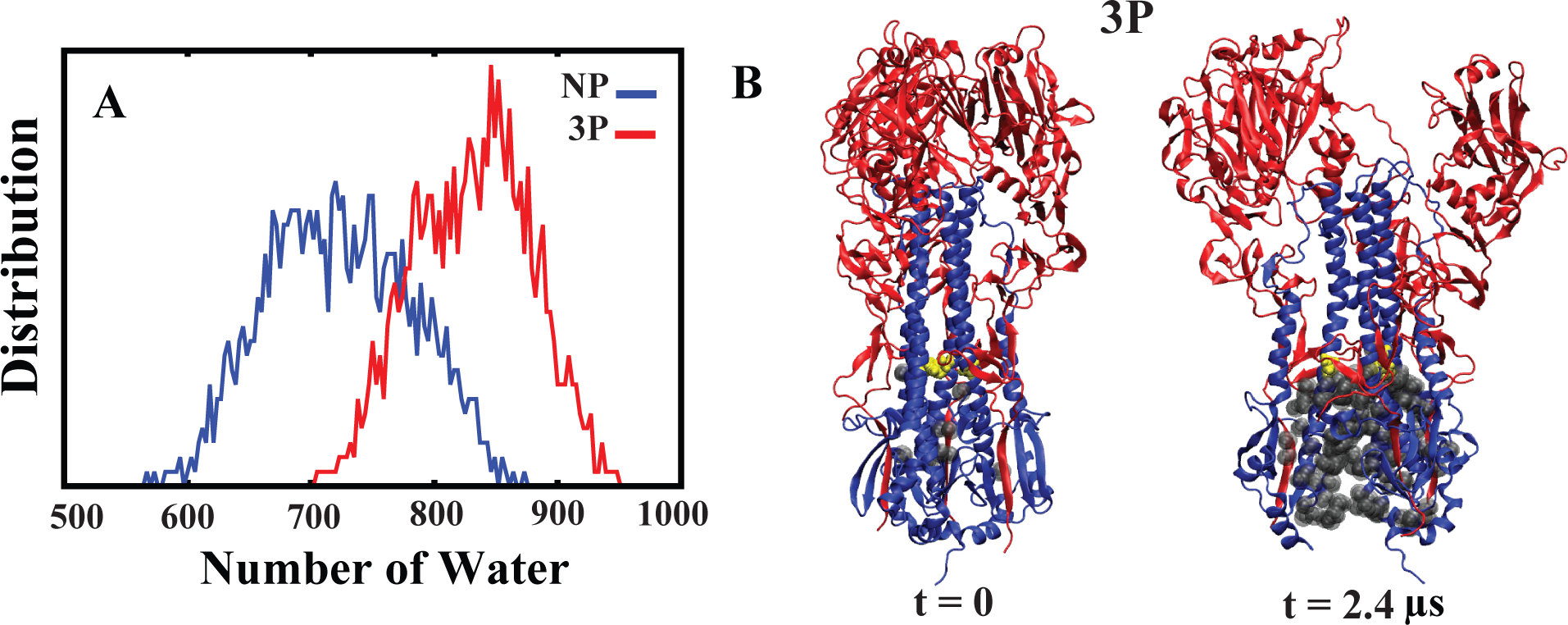
Water count probability distribution. (A) Probability distribution of the number of water molecules among three S4 helices during the final half of simulations (from 1.2 to 2.4 *µs*), comparing non-protonated (NP) and fully-protonated (3P) systems, based on three independent sets of simulations per condition. (B) Cartoon representation of the 3P system, showing the waters within the three S4 helices in the first and last frames of the simulation. Protonated histidines and water molecules are shown in yellow and gray, respectively.

**Fig. 5.**
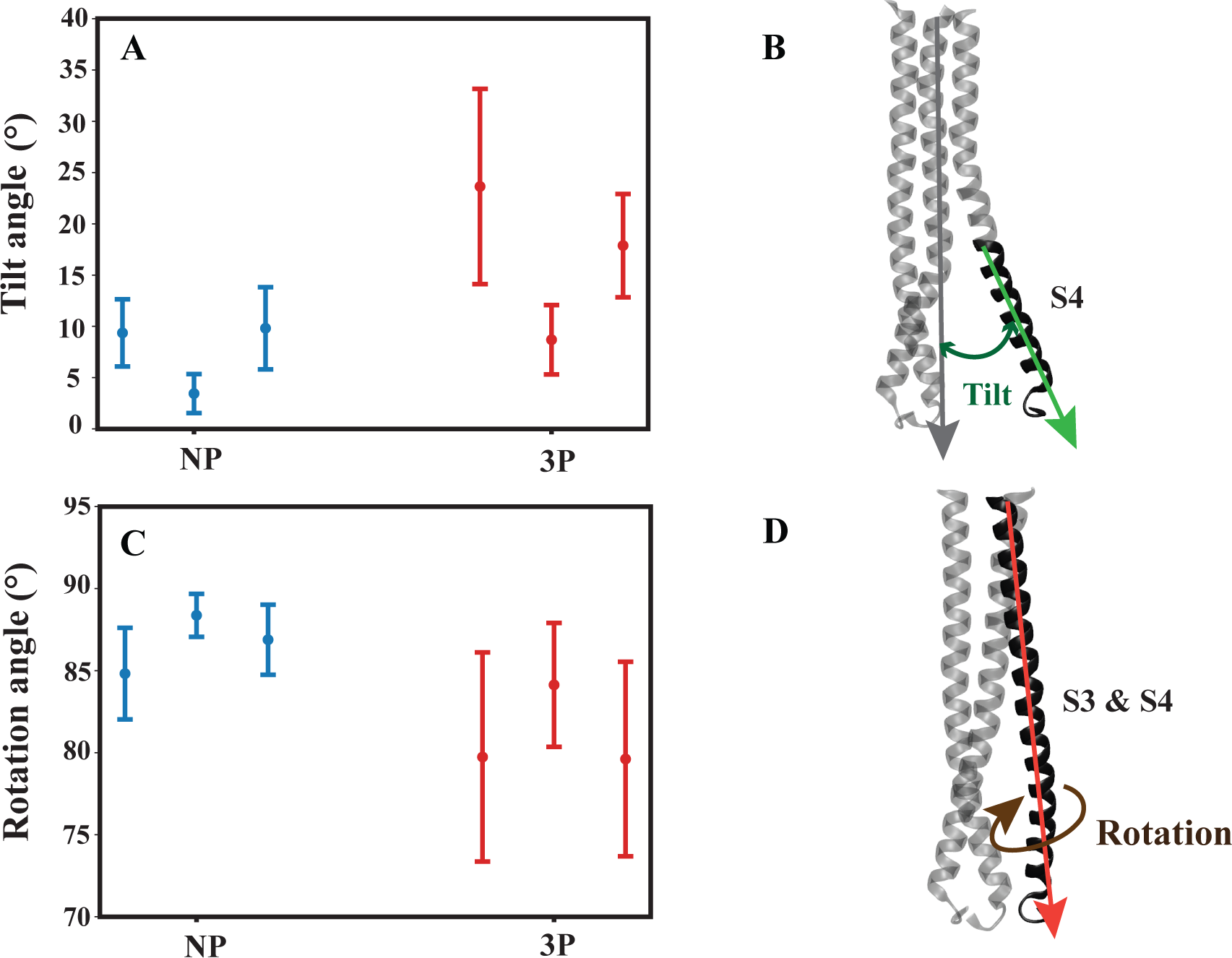
Average tilt and rotation angles. (A) The average tilt angle of the S4 helices for the last 1.2 *µs* of simulations in both non-protonated (NP, blue) and fully-protonated systems (3P, red), each repeated three times. (B) Definition of S4 tilt angle that is measured with respect to the entire long helices of HA2. (C) The average rotation angle of the long helices, including S3 and S4, for the last 1.2 *µs* in both non-protonated (NP, blue) and fully-protonated systems (3P, red), each repeated three times. (D) Definition of long helix rotation. The rotation of each monomer is calculated with respect to its crystal structure.

We found clear rotational differences in the side chain of the conserved histidine (His106_2_) between protonated and non-protonated systems when comparing the structural properties, with a special emphasis on conserved histidine residue located in the hinge region. We analyzed the angle of the long helix in all systems, combining both S3 and S4, to illustrate this rotation. The average initial angle for the 3P systems began at about 84 degrees, as shown in the figure (Fig. S3), and gradually reduced during the simulation, indicating a counterclockwise rotation. On the other hand, the angle increased or stayed the same in the NP systems, suggesting a rotation in a clockwise direction and inwards-facing conformation (Fig. S3). Additionally, the plot (Fig. 5B) depicting the average rotation angle of the 3P systems, considering both S3 and S4 domains across the three repeats, exhibited a reduction compared to the NP systems. The data points displayed a more scattered distribution in the 3P systems, indicating increased variability in rotational dynamics within these systems. According to our findings on tilt angle and rotational dynamics, it is evident that the protonation state plays a crucial role in shaping the conformational behavior of the S4 helix and the rotational dynamics of the conserved histidine residue (His106_2_).

In addition to our findings, relevant literature^62^ further supports the significance of our observations. A recent study focusing on influenza hemagglutinin (HA) highlighted the elusive nature of intermediate structures in HA conformational dynamics. This study utilized cryo-electron microscopy (cryo-EM) to identify two distinct low pH intermediate conformations of HA, revealing notable conformational changes in the central helices (S3 and S4). Specifically, these helices underwent an anticlockwise rotation of 9.5*^◦^* and a shift of 15 Å. In particular, the most significant conformational changes were observed in the helices corresponding to helix Ds, which aligns with our identification of S4 in our systems. This convergence of findings strengthens the relevance of our study and underscores the importance of understanding the dynamic behavior of HA in response to protonation states.^62^

### Investigating protonation effects on helix rotation in partially and fullyprotonated systems

In order to learn more about the underlying reasons behind this observed rotation, additional analyses such as salt bridge and hydrogen bond were conducted. The salt bridge analysis unveiled a significant interaction between His106_2_ from HA2 and its neighboring HA1 (Asp31_1_) located on 30-loop region, this bond appear to be absent in the NP systems (Fig. 6A-D). This finding provides insight into the potential role of protonation in stabilizing key intermolecular interactions, thereby influencing the rotational dynamics observed within the system.

**Fig. 6.**
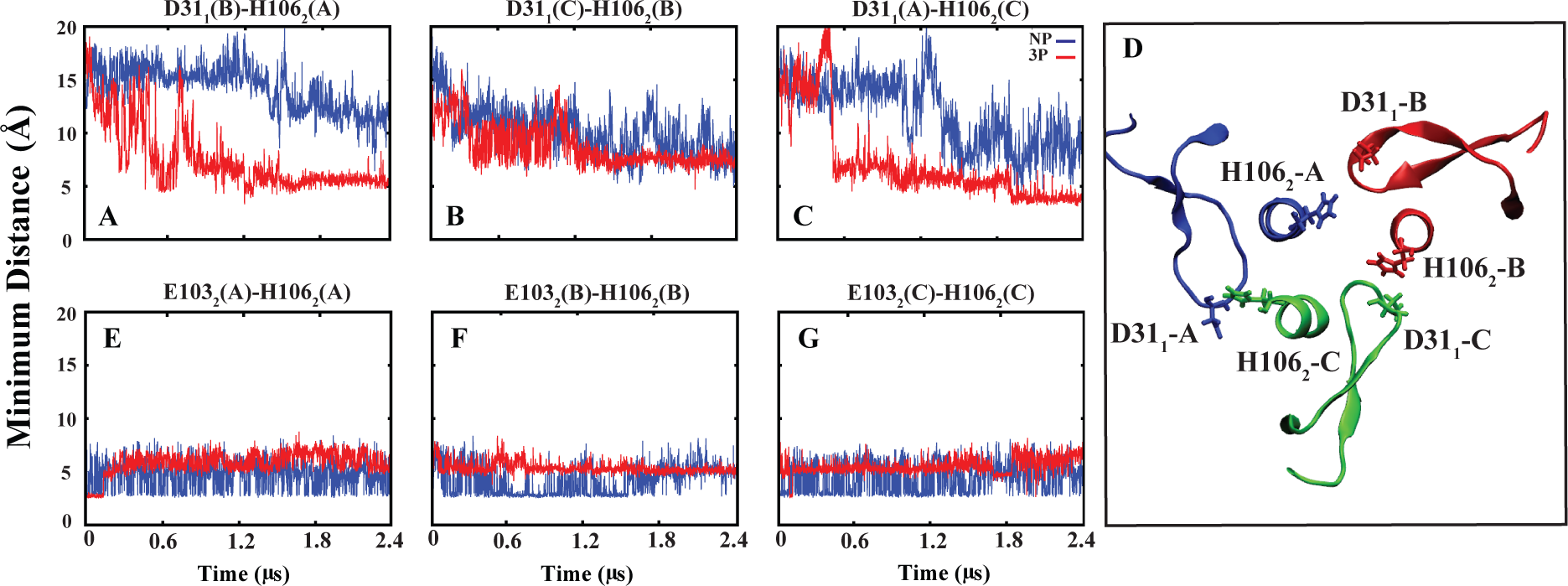
(A-C) Minimum distance between His106_2_ (HA2) and its neighboring Asp31_1_ (HA1). (D) Graphical representation of H106 and D31 in different monomer. (E-G) Minimum distance between His106_2_ and Glu103_2_, both located on the same monomer as HA2.

Measuring the minimum distance between Asp31_1_ and His106_2_ in all systems revealed interesting finding (Fig. S4). In the NP systems, this distance consistently remains greater than 5 Å, indicating the lack of a salt bridge formation. However, in the 3P systems, this salt bridge was observed in almost all simulation repeats, characterized by a significant decrease in the minimum distance between ASP31_1_ and His106_2_. Interestingly, analysis of partially protonated systems with 1 or 2 protonations (1P, 2P) displayed intermediate behavior between NP and 3P, further supporting the role of protonation states in modulating salt bridge formation and subsequent conformational dynamics (Fig. S4).

An investigation by Benton et al.(2020)^22^ employed cryo-EM of the HA ectodomain to explore structural transitions between HAs at neutral pH and fusion pH. Their study shed light on the concerted rearrangements of HA1 and HA2, particularly focusing on the 30-loop (HA1 residues 22–37). They found that some of the interaction of the 30-loop with the long helix of HA2 was similar in states I (neutral) and IV (extended HA2). In state IV, changes in side chains were observed, associated with the relocation of the short helix of HA2. Notably, in this state, the 30-loop formed interactions with His106_2_ and Gln105_2_ at the site of the 180*^◦^* turn in the fusion-pH structure, facilitating potential interactions between Thr30_1_ with

Gln105_2_ and His106_2_ of the adjacent HA2 chain. Additionally, the strictly conserved Asn104_2_ was observed to form hydrogen bonds with loop residue Lys27_1_ and Lys315_1_ of HA1, as well as HA2 residue Gln105_2_, suggesting potential roles for the 30-loop in the refolding process. In our own research, we have observed similar trends in the hydrogen bond occupancy between Asn104_2_ and Lys27_1_ and Lys315_1_ across different protonation states (occupancy around 70% for all systems such as protonated and non-protonated). However, our findings also highlight distinct interactions involving His106_2_, particularly with neighboring HA1 residue Asp31_1_ (30-loop) in our protonated systems (Fig. 6A-C).

Furthermore, in 3P systems, we observed that His106_2_ loses its interaction with Glu103_2_ of the same monomer HA2, whereas this interaction is more prominent in NP systems (Fig. 6E-G). The minimum distance calculation between His106_2_ and Glu103_2_ illustrates that while this interaction was evident in the NP systems and in some cases of partially protonated systems (1P and 2P), it failed to form or was disrupted at the onset of simulations in 3P systems (Fig. S5). Interestingly, it appears that this salt bridge predominantly forms during equilibration states and specially in monomers lacking protonation. These observations providing additional insights into the conformational dynamics of HA at different pH conditions.

In a study by Milder et al.(2022),^63^ the authors investigated the stabilization of prefusion HAs and their conformational impact through mutation of residues in the pH-sensitive region of HA2. By determining the X-ray crystal structure of the stabilized apo ectodomain of group 2 H3-HK68, they found that the mutations caused three histidines (His106_2_) to flip toward the trimer’s threefold axis, forming hydrogen bonds with Asp109_2_. Additionally, Ile29_1_ from the 30-loop was observed to lock the histidine (His106_2_) in the inward-facing conformation by shifting toward the threefold axis.

### The enhanced probability of fusion peptide release in protonated systems

An interesting aspect observed in our study revolves around the release of the FP within these systems. Despite the lack of direct observation of this event, strong evidence points to a significant difference in the stability of the FP between systems that are protonated and those that are not. Interestingly, indications point towards decreased stability of the FP in systems with protonation (fully or partially) compared to non-protonated ones. This observation highlights the potential influence of protonation states on the dynamics and stability of critical viral fusion machinery, providing important insights into the interaction between molecular behavior and environmental conditions during viral entrance processes. The conformational dynamics of the FP across all systems were investigated utilizing Principal Component Analysis (PCA), achieved through a combination of trajectories and projection onto the first and second principal components. As a result, systems with protonation exhibited distinct positions on the PCA plots, demonstrating divergence even within their non-protonated segments (Fig. 7).

**Fig. 7.**
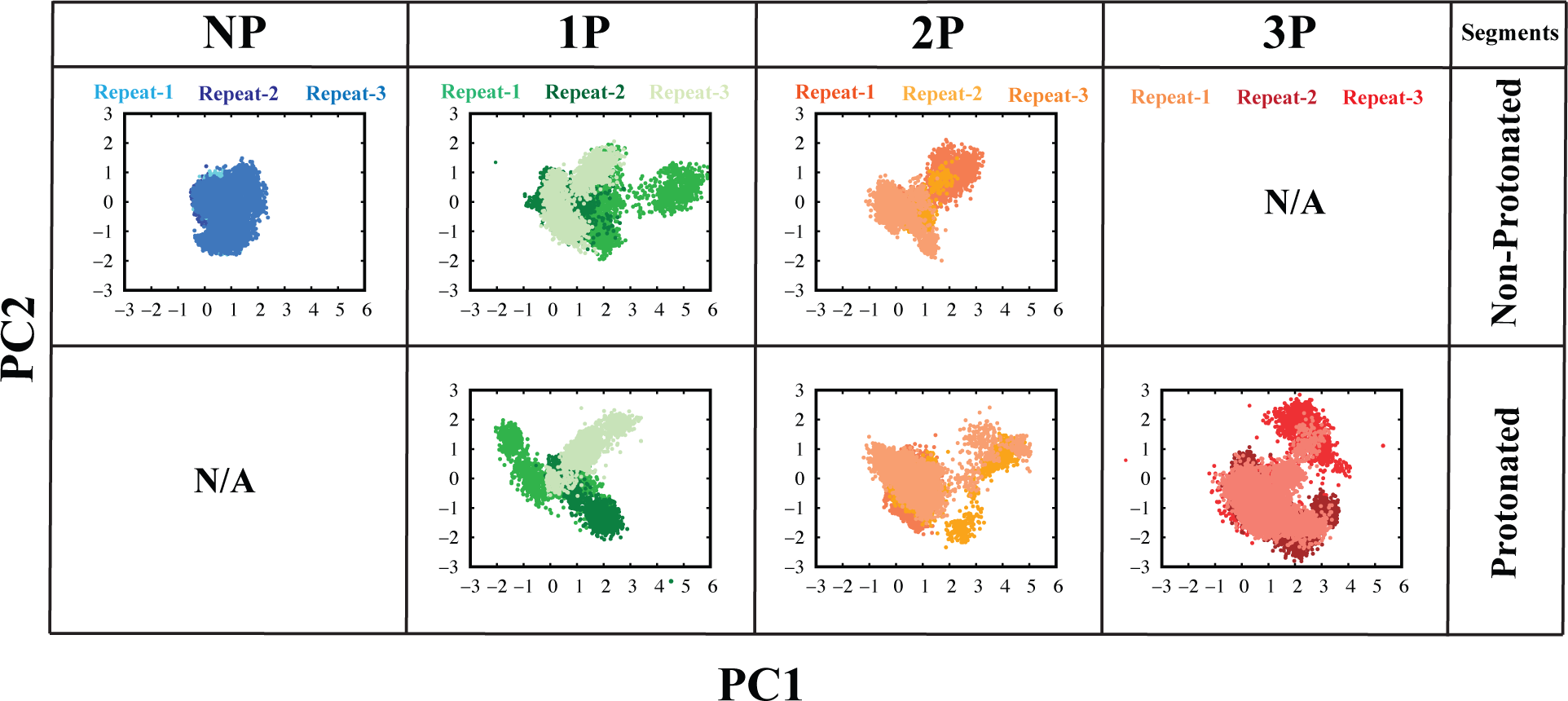
Projections of principal components (PCs) 1 and 2 depict the conformational landscape of the fusion peptide across non-protonated (NP), partially protonated (1P, 2P), and fully-protonated (3P) systems, as revealed by PCA. The first row represents non-protonated segments in each system, while the second row displays protonated segments. Each color corresponds to a specific system.

The hydrogen bond analysis revealed interactions involving the FP residue Leu2_2_ and neighboring residues, particularly Ser113_2_ and Asp109_2_ of S4 (Fig. 8A-F). Interestingly, an inter-hydrogen bond interaction between Leu2_2_ and Ser113_2_ (Fig. 8A-C) displayed higher occupancy in the NP system compared to protonated ones, particularly in the 3P sysytem. Plotting the hydrogen bond interaction unveiled its stability in NP systems, contrasting with its breakdown in partially or fully-protonated systems (1P, 2P, and 3P). Additionally, an intra-hydrogen bond interaction between Leu2_2_ and Asp109_2_ (Fig. 8D-F) within the hinge region was observed, albeit with decreased occupancy in protonated systems. Consistent with the previous observation, the plots illustrated the breaking of this bond in almost all protonated systems, while it remained stable in the NP systems (Fig. S6 & S7).

**Fig. 8.**
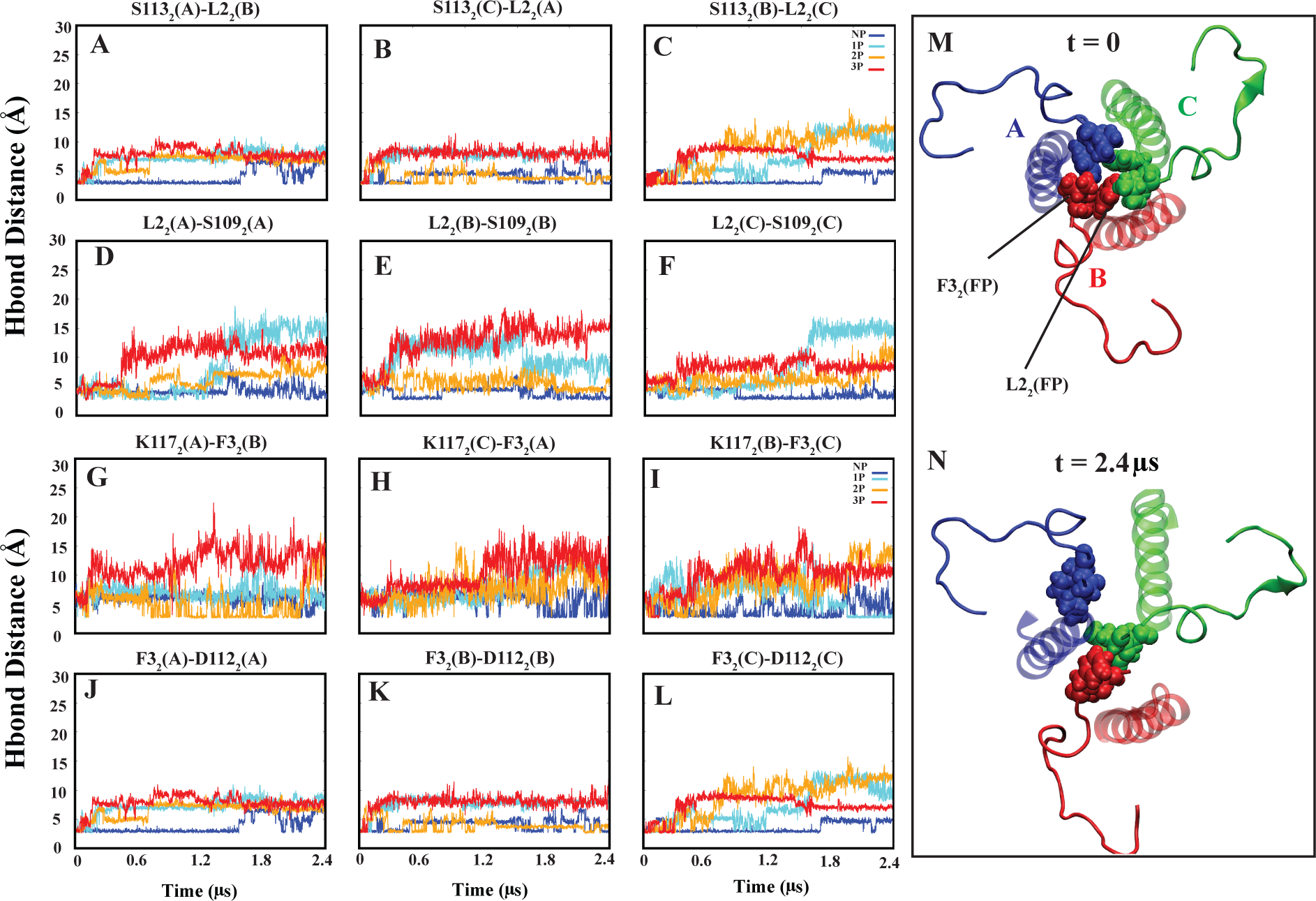
The inter- and intra-hydrogen bond distances between FP residues (Leu2_2_, Phe3_2_) and S4 residues (Ser113_2_, Asp109_2_, Lys117_2_, and Asp112_2_). (A-C and G-I) Denote inter-monomer hydrogen bonds between FP and S4 residues, indicating they are located in different monomers. (D-F and J-L) Represent intra-monomer hydrogen bonds between FP and S4 residues, signifying they are located within the same monomer.

Another notable intra-hydrogen bond interaction involves the Phe3_2_ residue of the FP with a neighboring residue of S4, namely Lys117_2_ (Fig. 8G-I). In particular, the average occupancy of this bond is nearly halved when comparing NP to 3P systems. The hydrogen bond plots provide further support for this observation, reflecting the trends observed in the average occupancy analysis. Similarly, an inter-hydrogen bond involving Phe3_2_ with another S4 residue, Asp112_2_ (Fig. 8J-L), displayed a decrease in average occupancy from NP to 3P systems. The H-bond interaction plots for FP (Phe3_2_) and S4 (Lys117_2_, and Asp112_2_) can be found in Supporting Information S8 and S9.

In a recent investigation conducted by Gao et al.(2020),^62^ the study focused on examining the structural intermediates during the low pH-induced transition of influenza hemagglutinin (HA). In the cryo-EM analysis of HA-antibody Fab complexes at neutral pH (pH 7.8), it was observed that the FP encircles the N-terminal fragment of HA1 (residues 9–19, HA1- N), forming a hydrogen bond with residue His17_1_ of HA1-N. Moreover, hydrophilic pockets are formed between the Helix Ds (S4s), allowing the hydrophobic distal ends of the FPs to penetrate deeply, creating a hydrophobic core consisting of residues Leu2_2_ and Phe3_2_). Gao et al. employed cryo-EM and 3D classification techniques to characterize the structures of HA in low pH-induced intermediate states. Their findings indicate that exposure to low pH triggers significant conformational changes in HA, leading to the release of the fusion peptide.

### Partial dissociation of HA1

Several studies have demonstrated that low pH induces the initial conformational change in HA by inducing partial dissociation or detachment of HA1 monomers.^22,27,64–71^ In our study, we observed that protonation similarly affects HA1 detachment. It triggers the formation of a cavity between HA1 and HA2, allowing water to penetrate and interact with the HA2 domain that was previously protected. This interaction potentially enhances the flexibility of the HA2 domain. To demonstrate these changes, we conducted a comprehensive analysis of the number of water molecules between the HA1 and HA2 domains in both NP and 3P systems. Our findings reveal that the number of water molecules in the 3P systems is higher compared to the NP systems (Fig. 9A & B), and the time dependent calculation of the water molecules is shown in Figure. S10. We further investigated the movement of HA1 by calculating the center of mass distance between its head (RBD & VE) and tail (F*^′^*) (Fig. 9 C & D) to assess the impact of protonation. The supporting information (Fig. S11) provides the time-dependent center of mass distance calculation. The center of mass distance between head and tail of the HA1 (Fig. 9C) indicate that the 3P systems exhibit slightly greater movement compared to the NP systems. These results indicate that protonation potentially influences the movement of HA1 and affects the hydration dynamics between HA1 and HA2 domains.

**Fig. 9.**
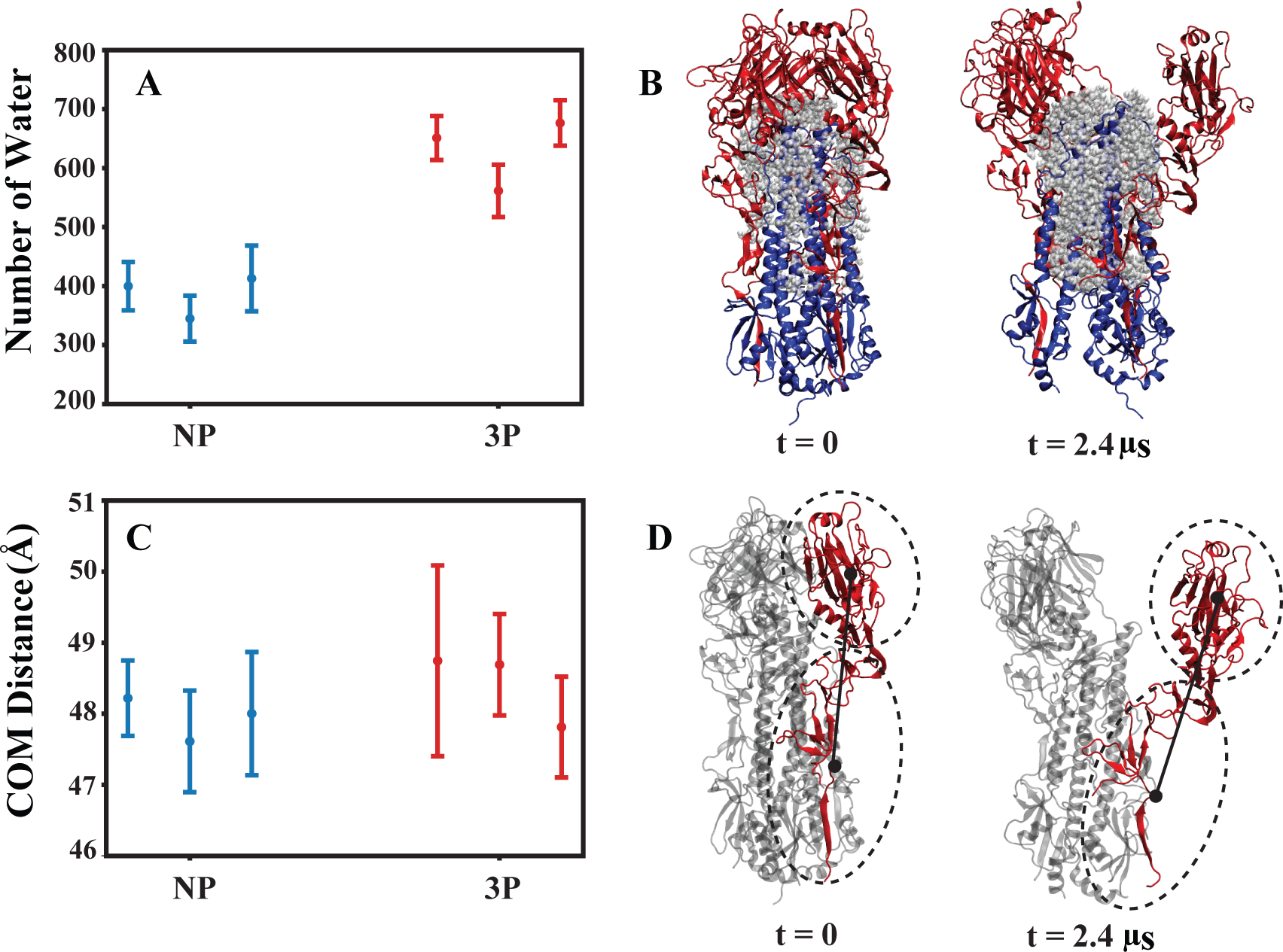
Average and standard deviation of penetrated water molecules and center of mass distances within each HA1. (A) Number of water molecules between HA1 and HA2 in non-protonated (NP) and fully-protonated (3P) systems for each repeat, depicting the last 1.2 *µs* of the simulations.(B) Graphic representation of the first and last frames of 3P system, with water molecules surrounding HA1 (red) and HA2 (blue) depicted in gray. (C) Measurement of the distance between the center of mass of the head and tail of HA1 for each repeat, focusing on the last 1.2 *µs* of the simulations. (D) Molecular image showing the center of mass distance considered between the head and tail of each HA1.

Based on our hydrogen bond analysis, we identified three crucial hydrogen bonds that exhibit significant differences between 3P and NP systems (Table. 1). The table presents the actual occupancy values for these hydrogen bonds in each system. Here, we summarize the average occupancies for clarity. The first bond, between His17_1_ and His18_1_ located on HA1, shows an average occupancy of 60% in NP systems, decreasing to 40% in 3P systems. The second hydrogen bond form between residues located on the 30-loop of HA1, Glu35_1_ and Gly23_1_, has an average interaction occupancy of 35% in NP systems, which decreases to 20% in 3P systems. The third one is H-bond between Asn322_1_ and Val20_1_ of HA1 domain with average occupancy of 59% in NP that reduced to 28% in 3P systems.

**Table 1:** Occupancy (%) of inter-domain H-bonds of HA1.

| System | H17<sub>1</sub>- H18<sub>1</sub> (%) |  |  | G23<sub>1</sub>- E35<sub>1</sub> (%) |  |  | N322<sub>1</sub>- V20<sub>1</sub> (%) |  |  |
| --- | --- | --- | --- | --- | --- | --- | --- | --- | --- |
|  | rep.1 | rep.2 | rep.3 | rep.1 | rep.2 | rep.3 | rep.1 | rep.2 | rep.3 |
| **NP** | 65 | 59 | 54 | 35 | 37 | 31 | 54 | 61 | 60 |
| **3P** | 33 | 44 | 41 | 18 | 25 | 17 | 19 | 35 | 30 |

There is a study^36^ that highlights the importance of conserved histidine residues, particularly HA1 position 17 (His17_1_), in triggering acid-induced conformational changes in the influenza hemagglutinin (HA) protein. His17_1_ is one of the conserved residues in the H3 subgroup, and extensive mutagenesis studies on these residues of H3 subtype HA suggest that His17_1_ may play a significant role in triggering structural changes upon acidification. This implies that His17_1_ is involved in the pH-dependent conformational changes necessary for membrane fusion during viral entry.^36^

Both histidine and glutamate residues are capable of accepting protons on their side chains, and we hypothesized that the protonation of His106_2_ could lead to an increase in water molecules around these residues, resulting in decreased interaction between the residues and an increased probability of their protonation. This, in turn, may create a more acidic environment, induce conformational changes in the protein, and enhance the probability of membrane fusion. We conducted an analysis of the water molecules surrounding the side chains of His17_1_ and Glu35_1_ to assess if their presence affects the interaction of residues (His17_1_-His18_1_ and Glu35_1_-Gly23_1_). Our results revealed a significant increase in the number of water molecules around these residues in the last 1.2 microseconds of the simulation in the 3P systems compared to the NP systems (Fig. 10A & B). The time series data of water around side chain of these two residues can also be found in Supporting Information (Fig. S12).

**Fig. 10.**
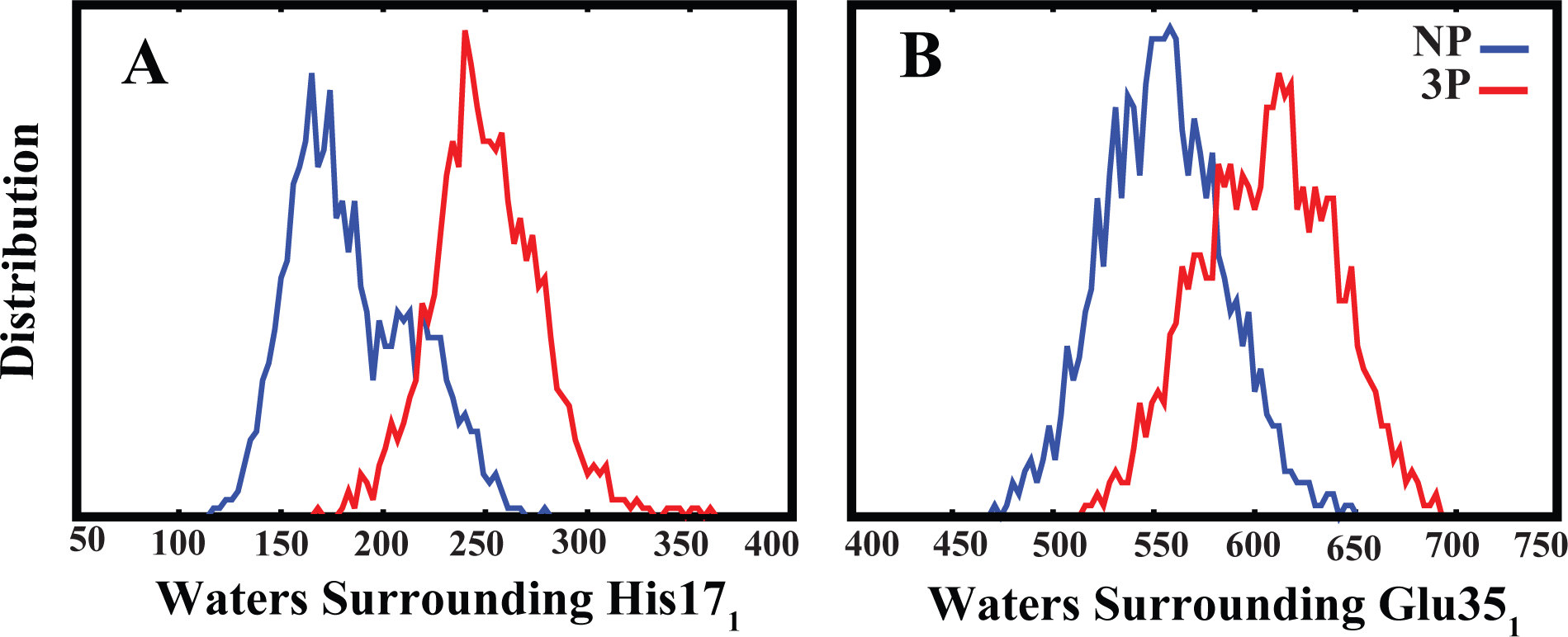
Comparing the number of water molecules surrounding the side chains of His17_1_ (A) and Glu35_1_ (B) in the HA1 domain, highlighting increased hydration in the fully-protonated systems (3P). Data from all three repeats for each system are included in these plots, focusing on the last 1.2 *µs* of the simulations.

## Conclusions

This work demonstrates that the protonation of His106_2_, a residue located in the hinge region of HA2, significantly impacts the fusion process of the HA protein. Our results indicate that protonating this specific residue induces notable conformational changes. These include decreased stability and the opening of the S4 helix, which contains the protonated histidine, along with an increase in water molecules between S4 helices. Additionally, His106_2_ residues exhibit an outward rotation in the protonated systems, whereas they maintain an inward-facing state in the NP systems. Furthermore, we observed initial signs of FP release and an increase in water molecules between HA1 and HA2. These changes trigger the conformational shifts in the protein that are necessary for facilitating the fusion process. Specifically, the conformational changes in HA1 include the center of mass distance between the head and tail of HA1 is altered in 3P systems compared to NP systems. The increased water penetration between HA1 and HA2 in the protonated systems disrupts hydrogen bonds, making more space around capable residues to attract protons and creating a more acidic environment that enhances the probability of membrane fusion. Our findings highlight the critical role of His106_2_ protonation in HA protein-mediated fusion. However, further studies are needed to deepen our understanding of protonation’s role in the fusion process. In the context of MD simulations, our research suggests that the protonation of His106_2_ can influence and potentially facilitate the fusion process.

## Acknowledgement

This research is supported by the National Science Foundation award CHE 1945465, the National Institutes of Health grants R15GM139140 and R35GM147423, and the Arkansas Biosciences Institute. This work also used the Extreme Science and Engineering Discovery Environment (XSEDE), which is supported by National Science Foundation Grant ACI-1548562. This work used XSEDE resources Comet and Stampede through allocation MCB150129. Anton 2 computer time was provided by the Pittsburgh Supercomputing Center (PSC) through Grant R01GM116961 from the National Institutes of Health. The Anton 2 machine at PSC was generously made available by D.E. Shaw Research.

## Supporting Information Available

This article contains Supporting Information.

