## Supporting Information for "Molecular dynamics investigation of the influenza hemagglutinin conformational changes in acidic pH"

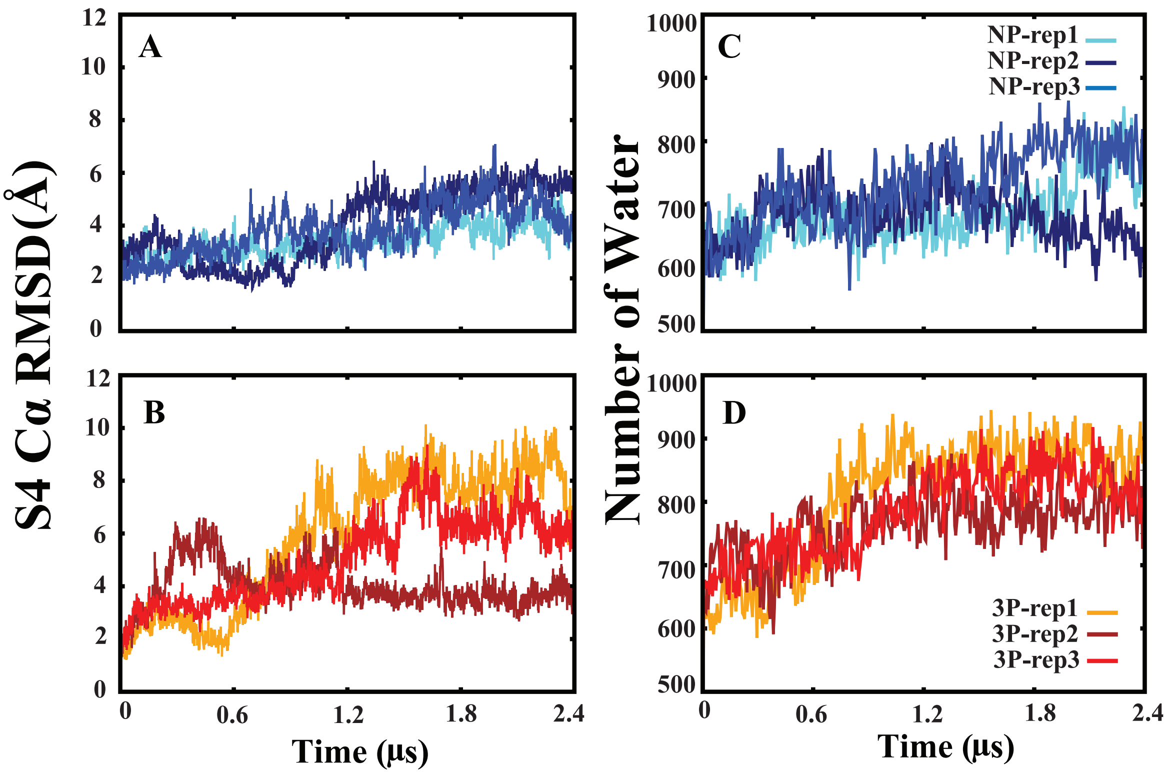


**Fig. S1. RMSD and water count analysis related to S4.** (A, B) The C$\alpha$ RMSD calculated for the S4 domains relative to the initial structure over the 2.4 $\mu$s simulation is plotted for both the non-protonated (NP) and fully-protonated (3P) simulations. (C, D) The number of water molecules within 3 Å of the three S4 helices of HA2, with each S4 from one monomer, was calculated during the 2.4 $\mu$s simulation. Each repeat is indicated with a different color.

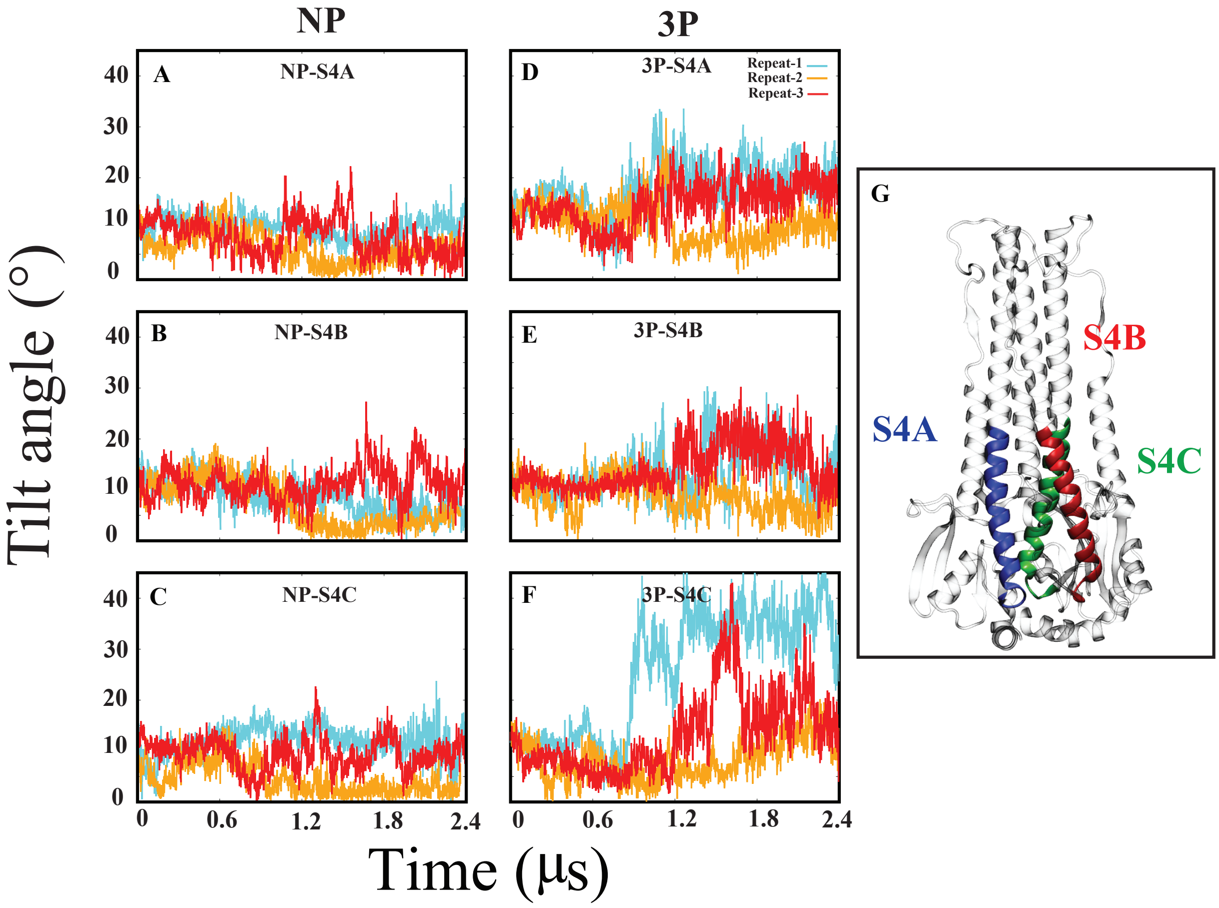


**Fig. S2. Tilt angle.** Time series of the tilt angle for S4 helix segments such as segment A (S4A), B (S4B), and C (S4C) in both non-protonated (NP) and fully-protonated (3P) systems. Each system undergoes three repetitions, with each simulation lasting 2.4 $\mu$s.

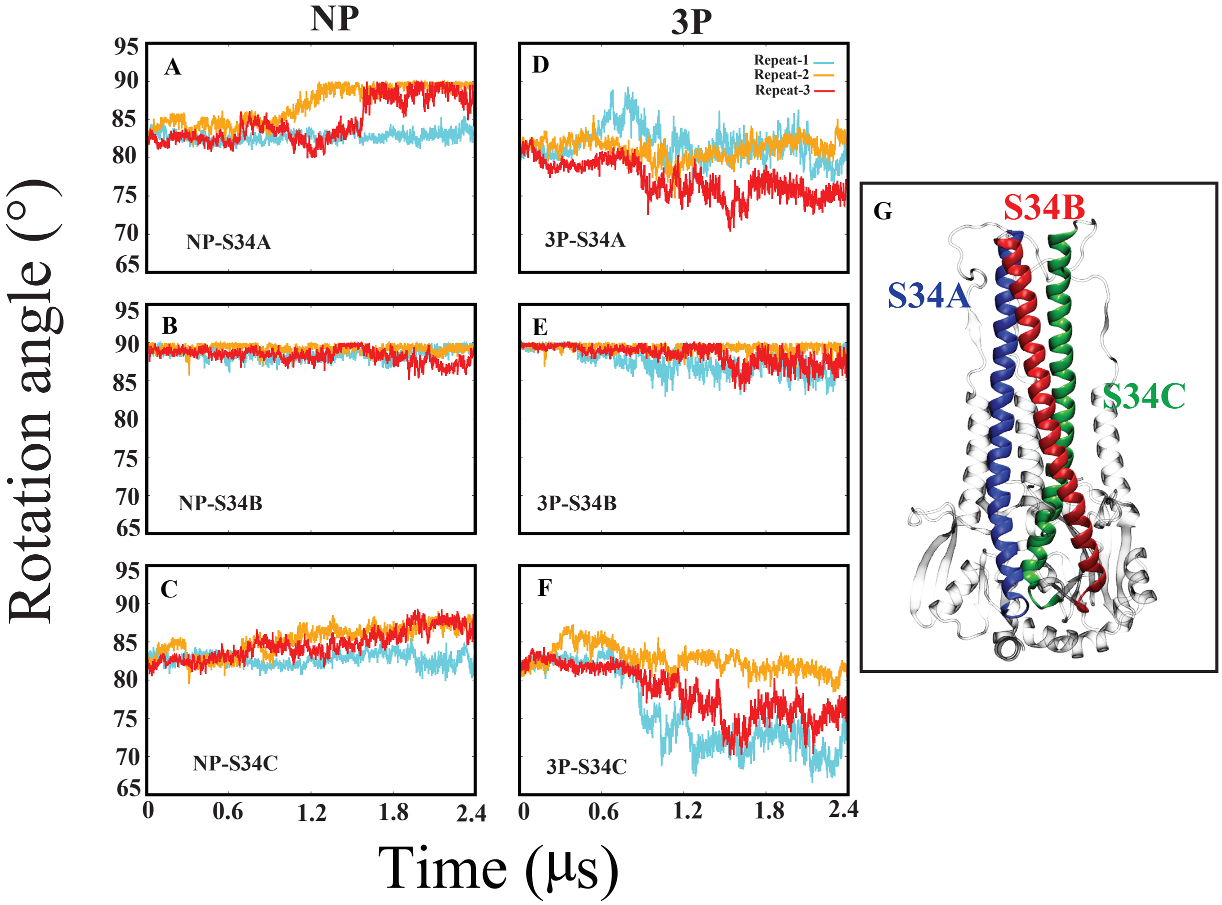


**Fig. S3. Rotation angle.** Time series of the rotation angle for the long helix of HA2, including S3 and S4 helices, segmented into A (S34A), B (S34B), and C (S34C), in both non-protonated (NP) and fully-protonated (3P) systems. Each system is repeated three times during the 2.4 $\mu$s simulation.

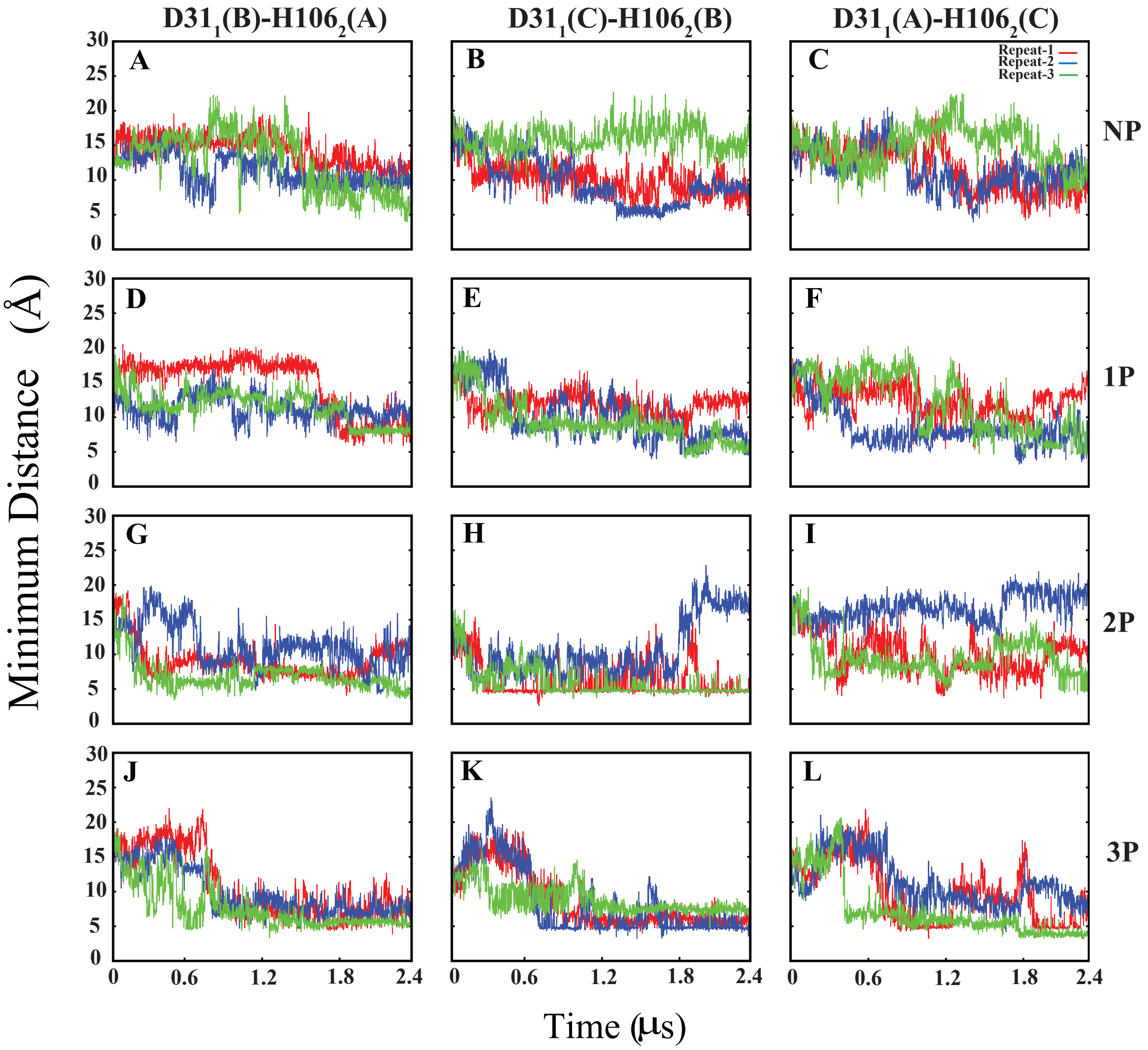


**Fig. S4. Minimum distance between H106$_2$ and neighboring D31$_1$ across protonation states.** Minimum distance between Asp31$_1$ and His106$_2$, located on HA1 and HA2, respectively. Rows represent different protonation states: non-protonated (NP), partially protonated (1P, 2P), and fully-protonated (3P). Each repeat is distinguished by a different color.

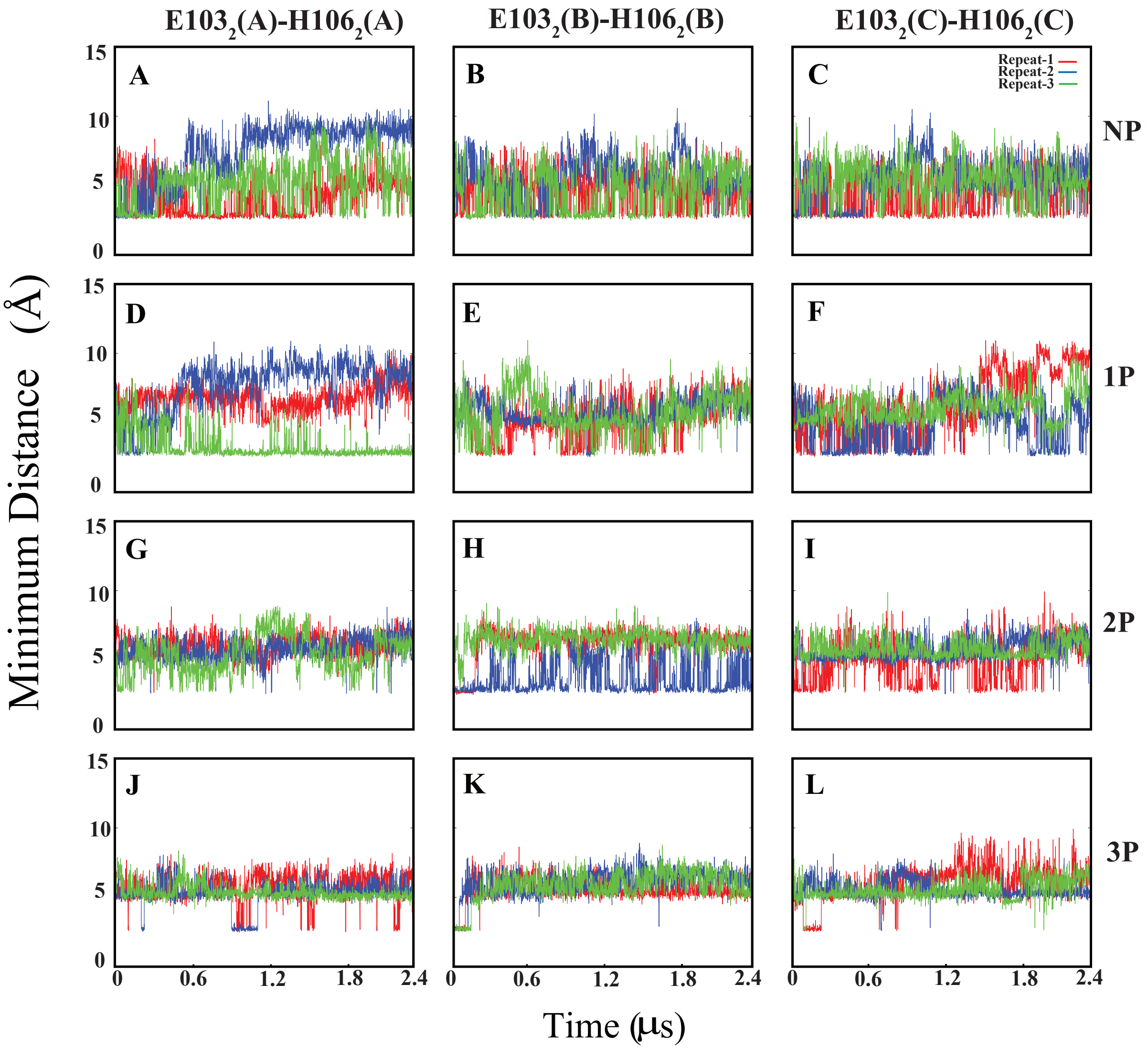


**Fig. S5. Minimum distance between Glu103$_2$ and His106$_2$ in the same monomer of HA2 across protonation states.** Minimum distance between Glu103$_2$ and His106$_2$, located on the same monomer of HA2, respectively. Rows represent different protonation states: non-protonated(NP), partially protonated (1P, 2P), and fully-protonated (3P). Each repeat is distinguished by a different color.

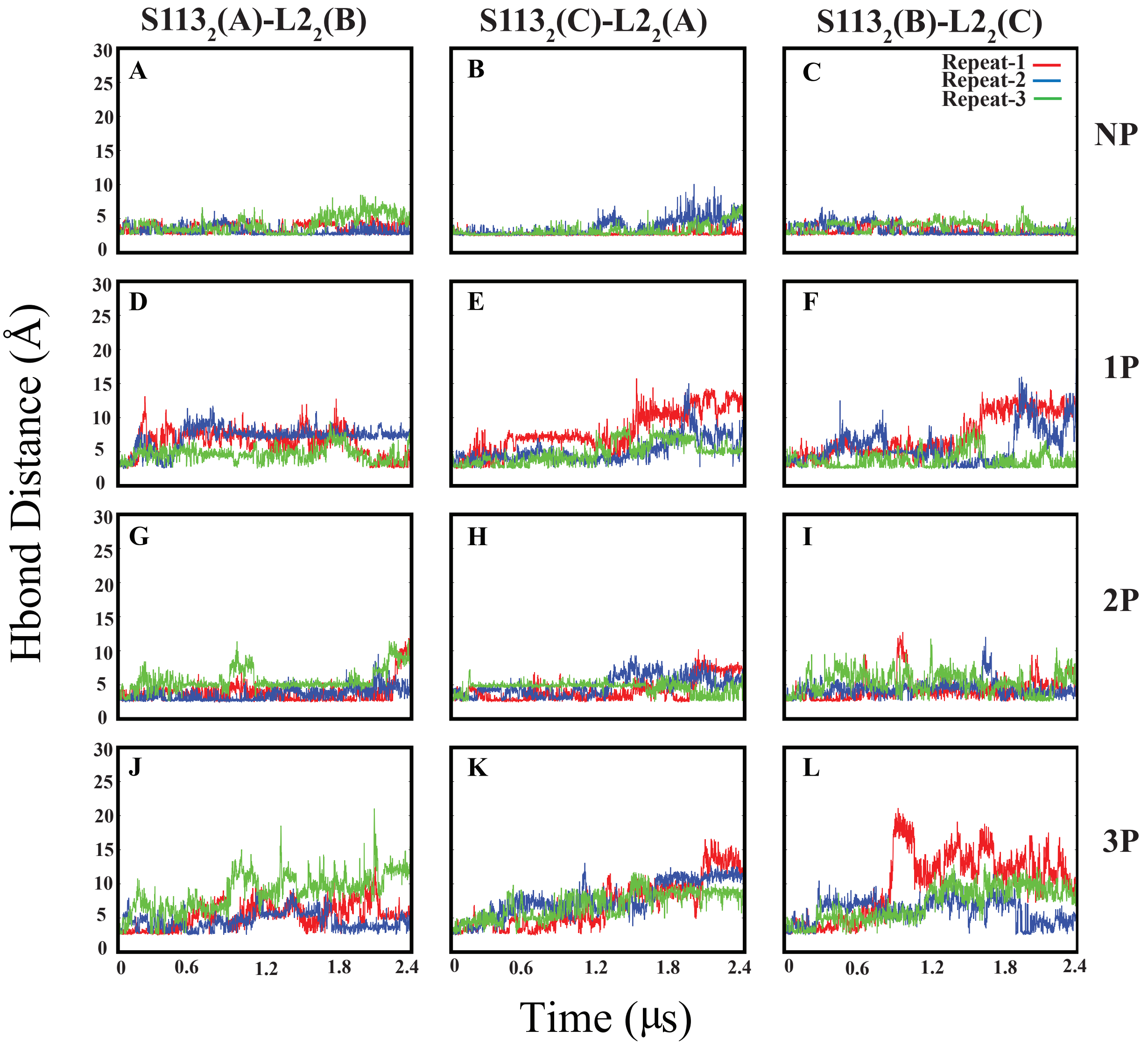


**Fig. S6. Inter-hydrogen bond distance between FP ($L2_2$) and S4 ($S113_2$) in different monomers of HA2 across protonation states.** Inter-hydrogen bond distance between $Leu2_2$ (FP) and $Ser113_2$ (S4), located on different monomers of HA2, during a 2.4 $\mu$s simulation. Rows represent different protonation states: non-protonated (NP), partially protonated (1P, 2P), and fully-protonated (3P). Each repeat is distinguished by a different color.

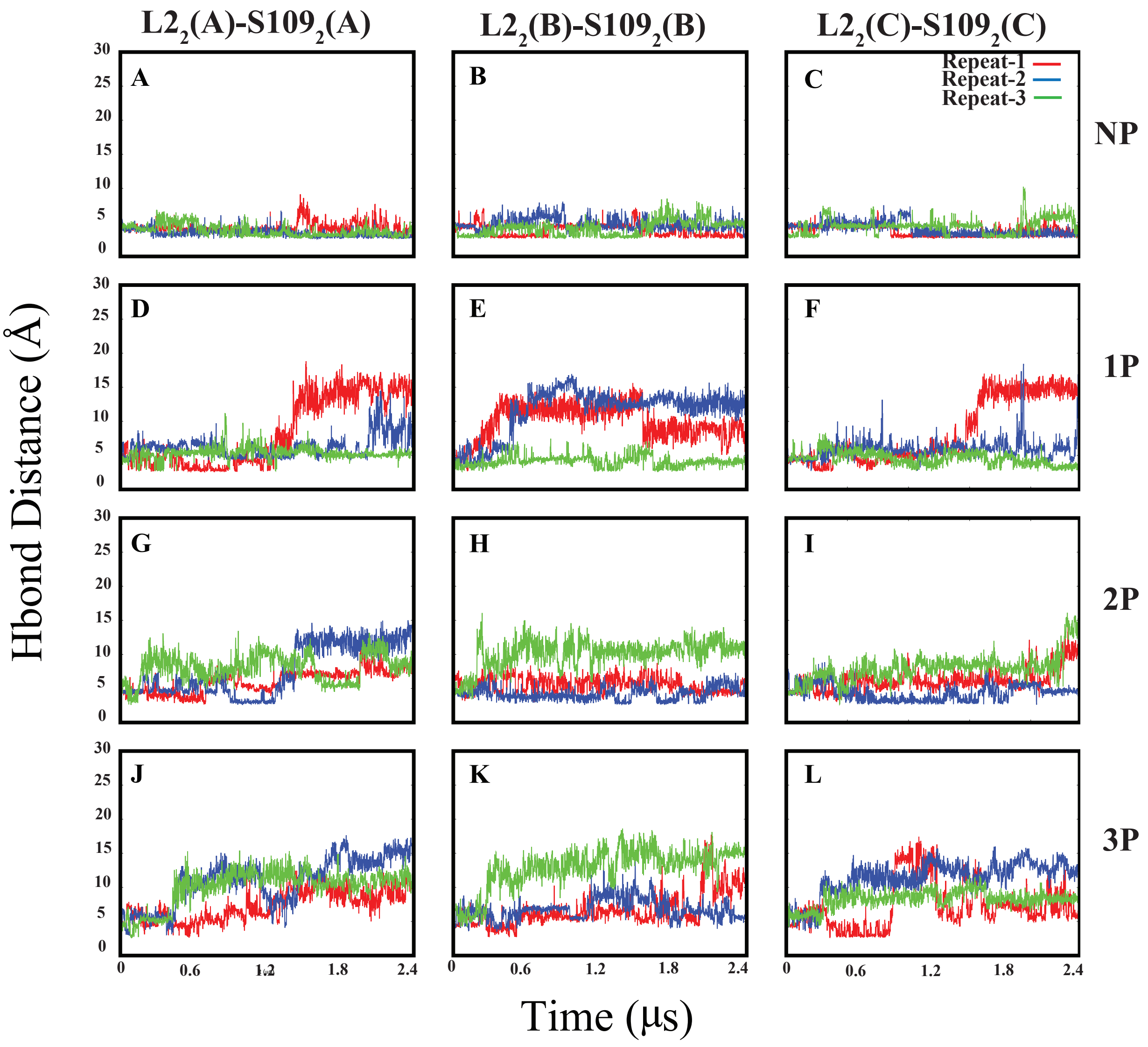


**Fig. S7. Intra-hydrogen bond distance between FP ($L2_2$) and S4 ($S109_2$) in the same monomers of HA2 across protonation states.** Intra-hydrogen bond distance between $Leu2_2$ (FP) and $Ser109_2$ (S4), located on the same monomers of HA2, during a 2.4 $\mu$s simulation. Rows represent different protonation states: non-protonated (NP), partially protonated (1P, 2P), and fully-protonated (3P). Each repeat is distinguished by a different color.

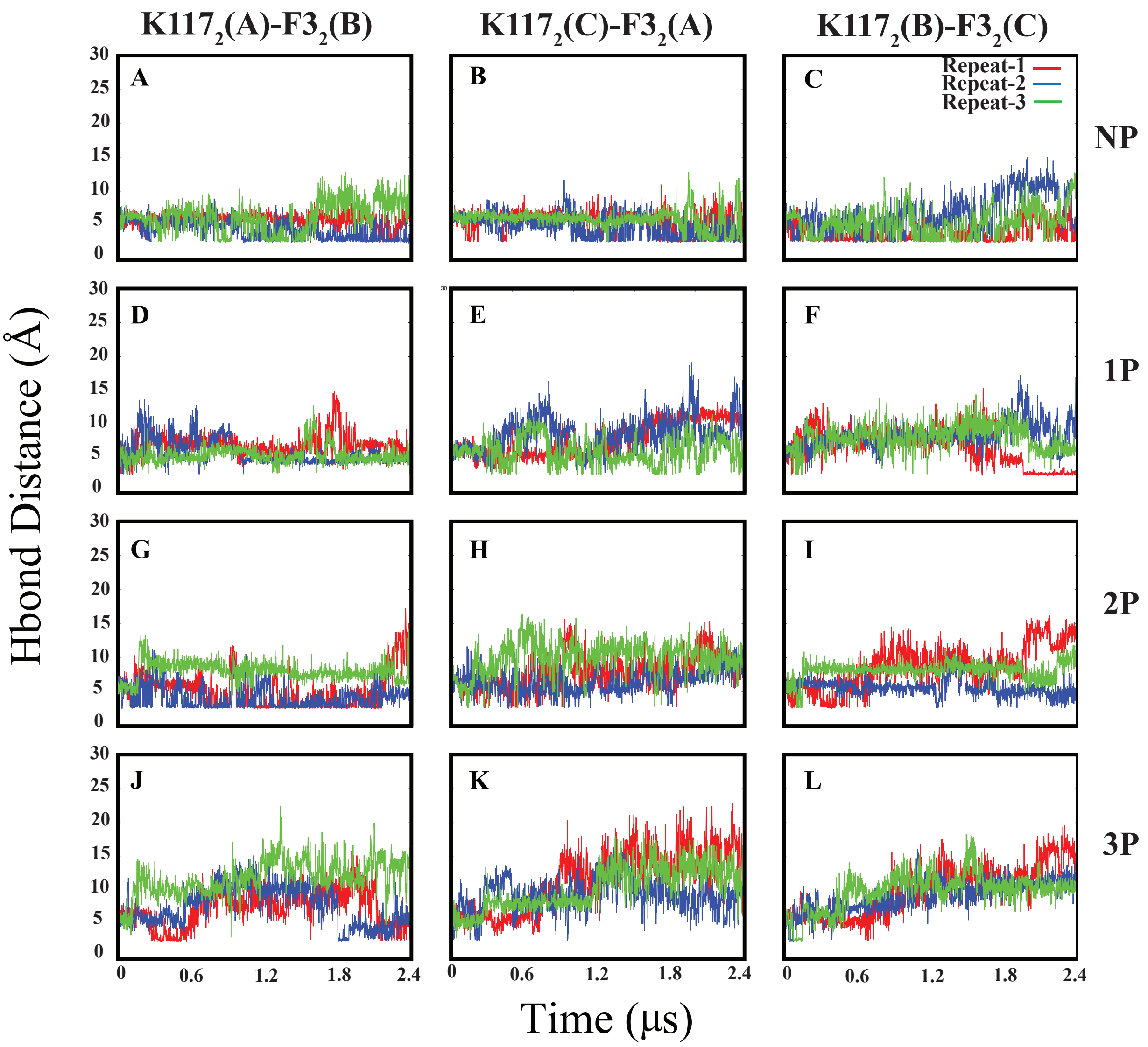


**Fig. S8. Inter-monomer hydrogen bond distance between FP ($F3_2$) and S4 ($K117_2$) in HA2 across protonation states.** Inter-monomer hydrogen bond distance between $Phe2_2$ (FP) and $Lys117_2$ (S4) in HA2, observed during a 2.4 $\mu$s simulation. Protonation states include non-protonated (NP), partially protonated (1P, 2P), and fully-protonated (3P), with each repeat distinguished by a different color.

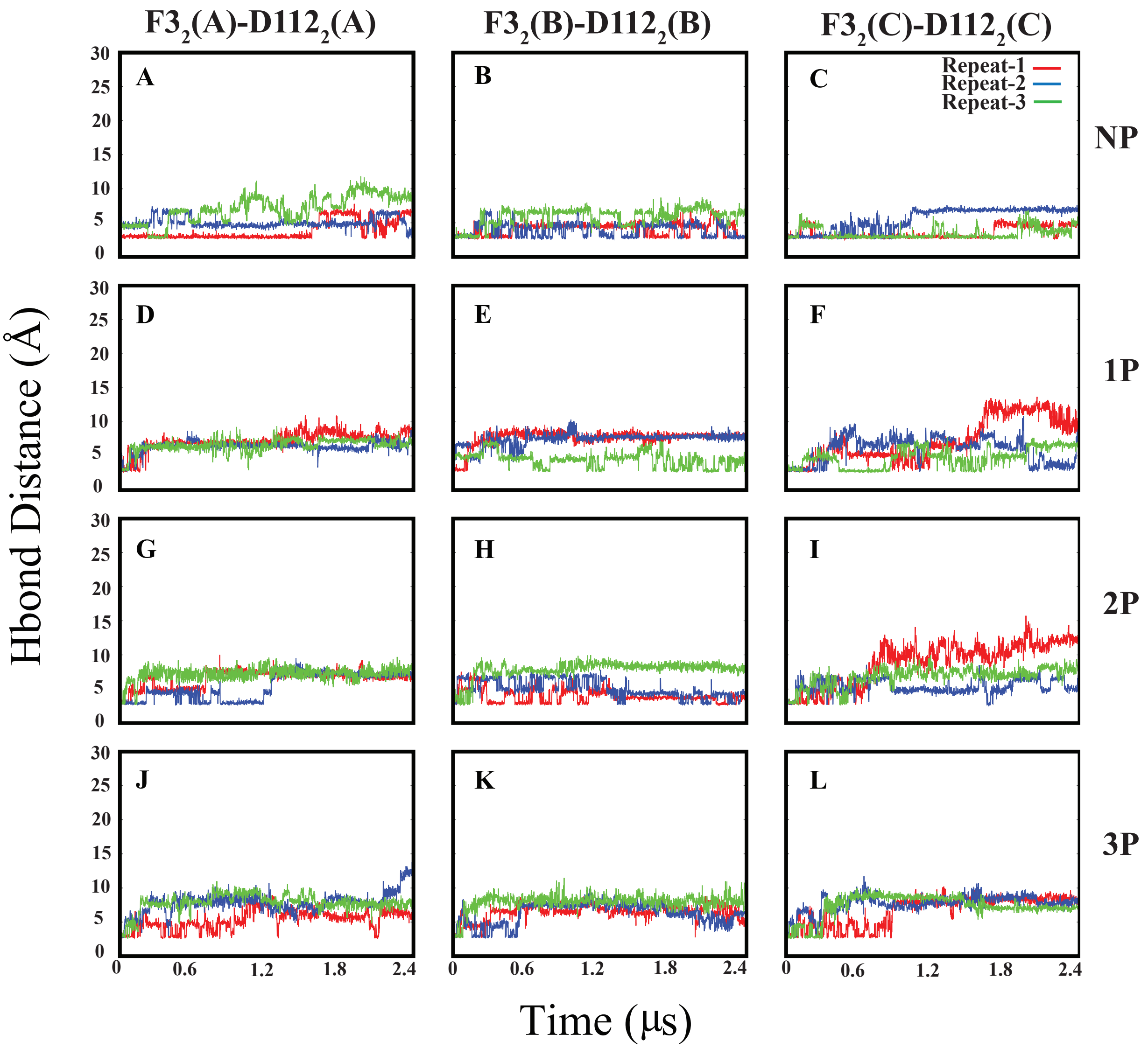


**Fig. S9. Intra-monomer hydrogen bond distance between FP ($F3_2$) and S4 ($D112_2$) in HA2 across protonation states.** Intra-hydrogen bond distance between $Phe3_2$ (FP) and $Asp112_2$ (S4), located on the same monomers of HA2, during a 2.4 $\mu$s simulation. Each row indicates different protonation states such as NP, 1P, 2P, and 3P, with each repeat distinguished by a different color.

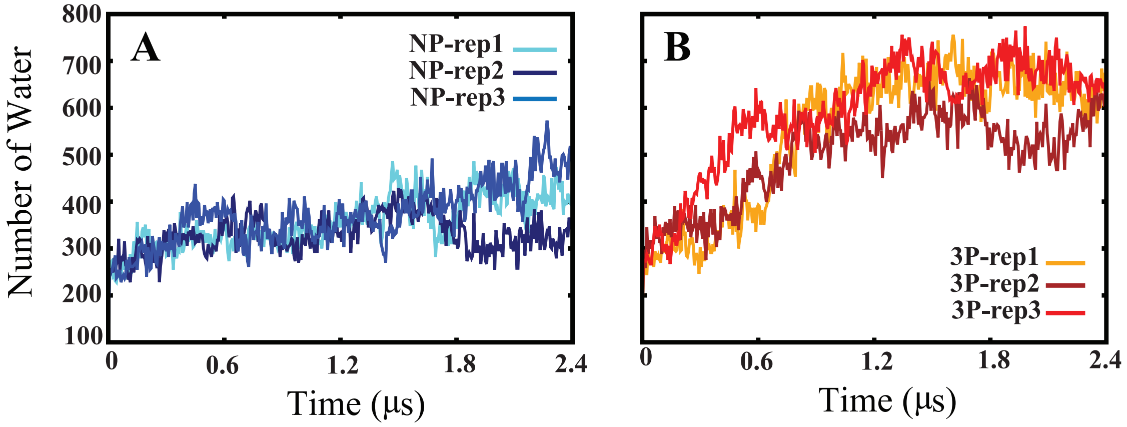


**Fig. S10. Water molecules between HA1 and HA2.** The number of water molecules calculated between HA1 and HA2 during the simulations in non-protonated (NP) and fully-protonated (3P) systems. Each repeat is indicated by a different color.

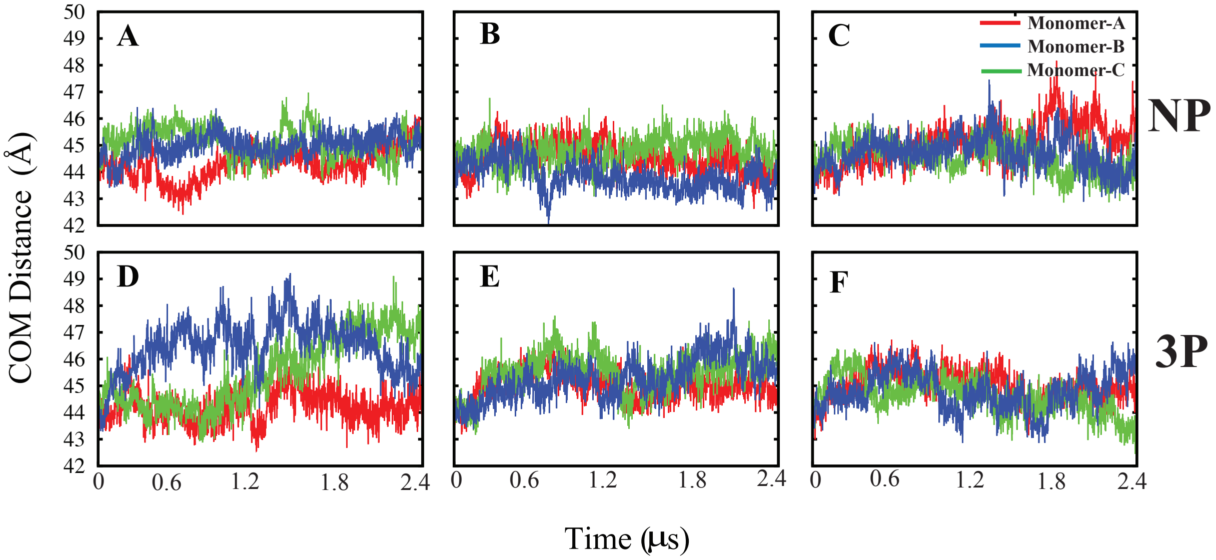


**Fig. S11. Center of mass distance between head and tail of the HA1 domain.** The center of mass distance between the head and tail of the HA1 domain during the simulations, with each monomer indicated by a different color.

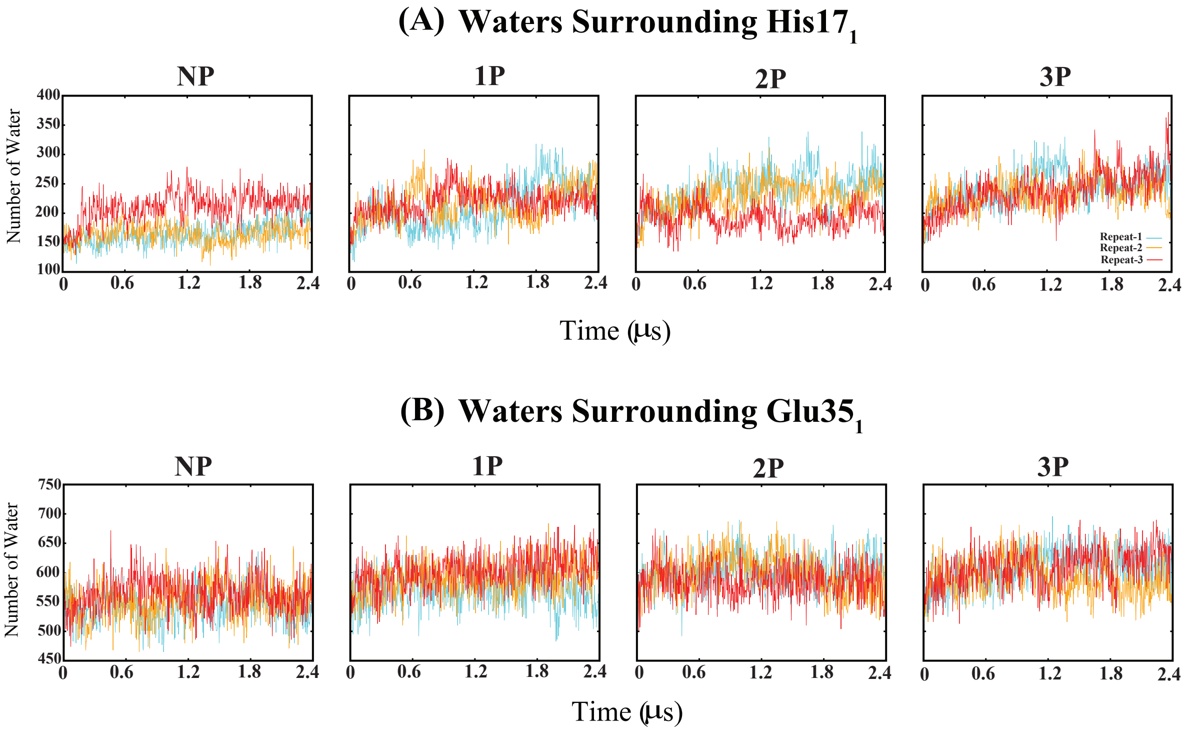


**Fig. S12. Water molecules around side chains in various protonation states.** The number of waters around the side chains of (A) residue His17$_1$ and (B) Glu35$_1$ calculated in non-protonated (NP), partially protonated (1P, 2P), and fully-protonated (3P) systems.
